# PGP-UK: a research and citizen science hybrid project in support of personalized medicine

**DOI:** 10.1101/288829

**Authors:** PGP-UK Consortium, Stephan Beck, Alison M Berner, Graham Bignell, Maggie Bond, Martin J Callanan, Olga Chervova, Lucia Conde, Manuel Corpas, Simone Ecker, Hannah R Elliott, Silvana A Fioramonti, Adrienne M Flanagan, Ricarda Gaentzsch, David Graham, Deirdre Gribbin, José Afonso Guerra-Assunção, Rifat Hamoudi, Vincent Harding, Paul L Harrison, Javier Herrero, Jana Hofmann, Erica Jones, Saif Khan, Jane Kaye, Polly Kerr, Emanuele Libertini, Laura McCormack, Ismail Moghul, Nikolas Pontikos, Sharmini Rajanayagam, Kirti Rana, Momodou Semega-Janneh, Colin P Smith, Louise Strom, Sevgi Umur, Amy P Webster, Karen Wint, John N Wood

## Abstract

Molecular analyses such as whole-genome sequencing have become routine and are expected to be transformational for future healthcare and lifestyle decisions. Population-wide implementation of such analyses is, however, not without challenges, and multiple studies are ongoing to identify what these are and explore how they can be addressed. Defined as a research project, the Personal Genome Project UK (PGP-UK) is part of the global PGP network and focuses on open data sharing and citizen science to advance and accelerate personalized genomics and medicine. Here we report our findings on using an open consent recruitment protocol, active participant involvement, open access release of personal genome, methylome and transcriptome data and associated analyses, including 47 new variants predicted to affect gene function and innovative reports based on the analysis of genetic and epigenetic variants. For this pilot study, we recruited ten participants willing to actively engage as citizen scientists with the project. In addition, we introduce Genome Donation as a novel mechanism for openly sharing previously restricted data and discuss the first three donations received. Lastly, we present GenoME, a free, open-source educational app suitable for the lay public to allow exploration of personal genomes. Our findings demonstrate that citizen science-based approaches like PGP-UK have an important role to play in the public awareness, acceptance and implementation of genomics and personalized medicine.

## Background

The sequencing of the first human genome in 2001 [1, 2] catalysed a revolution in technology development, resulting in around one million human genomes having been sequenced to date at ever decreasing costs [3]. This still expanding effort is underpinned by a widespread consensus among researchers, clinicians and politicians that ‘omics’ in one form or another will transform biomedical research, healthcare and lifestyle decisions. For this transformation to happen successfully, the provision of choices that accommodate the differing needs and priorities of science and society are necessary. The clinical need is being addressed by efforts such as the Genomics England 100K Genome Project [4] and the US Precision Medicine Initiative [5] (recently renamed to ‘All of Us’) whilst the public’s desire for direct-to-consumer genetic testing is met by a growing number of companies [6]. However, little of the data from these sources are being made available for research under open access which, in the past, has been a driving force for discovery and tool development [7]. This important research need for unrestricted access to data was first recognised by the Human Genome Project and implemented in the ‘Bermuda Principles’ [8]. The concept proved highly successful and was developed further by personal genome projects such as PGP [9-12] and iPOP [13] and, more recently, has also been adopted by some medical genome projects like TXCRB [14] and MSSNG [11] the latter of which uses a variant of registered access [15].

Founded in 2013, PGP-UK was the first project in Europe to implement the open consent framework [16] pioneered by the Harvard PGP for participant recruitment and data release. Under this ethics approved framework, PGP-UK participants agree for their omics and associated trait, phenotype and health data to be deposited in public databases under open access. Despite the risks associated with sharing identifiable personal information, PGP-UK has received an enthusiastic response by prospective participants and even had to pause enrolment after more than 10,000 people registered interest within a month of launching the project. The rigorous enrolment procedure includes a single exam to document that the risks as well as the benefits of open data sharing have been understood by prospective participants and the first 1,000 have been allowed to fully enrol and consent.

Taking advantage of PGP-UK being a hybrid between a research and a citizen science project, we (the researchers and participants) describe here our initial findings from the pilot study of the first ten participants and the resulting variant reports. Specifically, this includes the description of variants identified in the participants’ genomes and methylomes as well as our interpretation relating to ancestry, predicted traits, self-reported phenotypes and environmental exposures. As examples of citizen science, which we define here as activity that encourages members of the public to participate in research by taking on the roles of both subject and scientist [17], we describe the first three genome donations received by PGP-UK and a first genome app (GenoME) developed as educational tool for the lay public to better understand personalized and medical genomics. Mobile apps have become the method of choice for the public to engage with complex information and processes such as navigation using global positioning systems, internet shopping/banking and a variety of educational and health-related activities [18]. The open nature of the PGP-UK data make them an attractive resource for investigating interactions between genomics, environmental exposures, health-related behaviours and outcomes in health and disease. For example, the MedSeq Project recently trialled the impact of whole-genome sequencing (WGS) on the primary care and outcomes of healthy adult patients and identified sample size as one of the limiting factors [19].

## Methods

### Ethics

The research conformed to the requirements of the Declaration of Helsinki, UK national laws and to UK regulatory requirements for medical research. All participants were informed, consented, subjected to an online entrance exam and enrolled as described on the PGP-UK sign-up web site (www.personalgenomes.org.uk/sign-up). The study was approved by the UCL Research Ethics Committee (ID Number 4700/001) and is subject to annual reviews and renewals.

### Genome Donations

Ethics approval for PGP-UK to receive genomes, exomes and genotypes (e.g. 23andMe) and associated data generated elsewhere was obtained from the UCL Research Ethics Committee through an amendment of ID Number 4700/001. Enrolment in PGP-UK is accepted as proof that prospective donors have been adequately informed and have understood the risks of holding and donating their genome and associated data. Equal to regular participants, donors agree for their data and associated reports to be made publicly available under open access by PGP-UK. Once a genome donation has been received, the data are processed and reports produced as for genomes generated by PGP-UK. Donors are also eligible to provide samples for the generation of additional data and reports as implemented here for 450K methylome analysis.

### Samples

Blood samples (2 x 4 ml) were taken by a medical doctor at the UCL Hospital using EDTA Vacutainers (Becton Dickinson). Saliva samples were collected using Oragene OG-500 self-sampling kits (DNA Genotek). All samples were processed and stored at the UCL/UCLH Biobank for Studying Health and Disease (http://www.ucl.ac.uk/human-tissue/hta-biobanks/UCL-Cancer-Institute) using HTA-approved standard operating procedures (SOPs).

### Data generation and analysis

**Whole-genome sequencing (WGS)** was subcontracted to the Kinghorn Centre for Clinical Genomics (Australia) and conducted on an Illumina HiSeq X platform. Illumina TruSeq Nano libraries were prepared according to SOPs and sequenced to an average depth of 30X. The sequenced reads were trimmed using TrimGalore (http://www.bioinformatics.babraham.ac.uk/projects/trim_galore/) and mapped against the hg19 (GRCh37) human reference genome using the BWA-MEM algorithm from BWA v0.7.12 [20]. After removing ambiguously mapped reads (MAPQ < 10) with SAMtools 1.2 [21] and marking duplicated reads with Picard 1.130 (http://broadinstitute.github.io/picard/), genomic variants were called following the Genome Analysis toolkit (GATK 3.4-46; https://software.broadinstitute.org/gatk/) best practices, which involves local realignment around indels, base quality score recalibration, variant calling using the GATK HaplotyeCaller, variant filtering using the variant quality scoring recalibration (VQSR) protocol, and genotype refinement for high-quality identification of individual genotypes. Additionally, variants of phenotypic interest identified from SNPedia [22] that were not called using the above pipeline due to being identical to the human reference genome (homozygous reference variants), were obtained by preselecting a list of phenotypically interesting variants and requesting the GATK HaplotypeCaller to emit genotypes on these chromosomal locations. The WGS data (FASTQ and BAM files) have been deposited in the European Nucleotide Archive (ENA) under accession number PRJEB24961. The variant files (VCFs) have been deposited in the European Variant Archive (EVA) under accession number PRJEB17529 and linked to the Global Alliance for Genomics and Health (GA4GH) Beacon project (https://www.ebi.ac.uk/eva/?GA4GH) under the same accession number.

**Whole-genome bisulfite sequencing (WGBS)** was subcontracted to the National Genomics Infrastructure Science for Life Laboratory (Sweden) and conducted on an Illumina HiSeq X platform. Bisulfite conversion and library preparation were carried out using a TruMethyl Whole Genome Kit v2.1 (Cambridge Epigenetix, now marketed by NuGEN) and libraries sequenced to an average depth of 15X. The resulting FASTQ files were analysed using GEMBS [23]. As reported previously [24], WGBS on the Illumina HiSeq X platform is not straightforward as the data are of inferior quality to those that can be obtained on other HiSeq or NovaSeq platforms. In our case, the average unique mapping quality was 63.86% for paired-end (PE) and 86.18% for single-end (SE, forward) reads as assessed with GEMBS [23]. The WGBS data have been deposited in ENA under accession number PRJEB24961.

**Genome-wide DNA methylation profiling** was conducted with Infinium HumanMethylation450 (450K) BeadChips (Illumina). Genomic DNA (500ng) was bisulfite-converted using an EZ DNA Methylation Kit (Zymo Research) and processed by UCL Genomics using SOPs for hybridisation to 450K BeadChips, single-nucleotide extension followed by immunohistochemistry staining using a Freedom EVO robot (Tecan) and imaging using an iScan Microarray Scanner (Illumina). The resulting data were quality controlled and analysed using the ChAMP [25, 26] and minfi [27] analysis pipelines. The 450K data have been deposited in ArrayExpress under accession number E-MTAB-5377.

**RNA sequencing (RNA-seq)** was carried out on RNA extracted from blood using both targeted and whole RNA-seq. For targeted RNA-seq, library preparation was carried out using AmpliSeq (Thermo Fisher Scientific). A barcoded cDNA library was first generated with SuperScript VILO cDNA Synthesis kit from 20 ng of total RNA treated with Turbo DNase (Thermo Fisher Scientific), followed by amplification using Ion AmpliSeq technology. Amplified cDNA libraries were QC-analysed using Agilent Bioanalyzer high sensitivity chips. Libraries were then diluted to 100 pM and pooled equally, with two individual samples per pool. Pooled libraries were amplified using emulsion PCR on Ion Torrent OneTouch2 instruments (OT2) following manufacturer’s instructions and then sequenced on an Ion Torrent Proton sequencing system, using Ion PI kit and chip version 2.

For whole RNA-seq, the libraries were prepared from 20 ng of total RNA with Illumina-compatible SENSE mRNA-Seq Library Prep Kit V2 (Lexogen, NH, USA) according to the manufacturer’s protocol. The resulting double-stranded library was purified and amplified (18

PCR cycles) prior to adding the adaptors and indexes. The final PCR product (sequencing library) was purified using SPRI (Solid Phase Reversible Immobilisation) beads followed by

library quality control check, quantified using Qubit fluorometer (Thermo Fisher Scientific) and QC-analysed on Bioanalyzer 2100 (Agilent) and further quantified by qPCR using KAPA library quantification kit for Illumina (Kapa Biosystems). The libraries were sequenced on HiSeq 4000 (Illumina) for 150bp paired-end chemistry according to manufacturer’s protocol. The average raw read per sample was 36,632,921 reads and the number of expressed transcripts per sample was 25,182.

The RNA-seq data have been deposited in ArrayExpress under accession number E-MTAB-6523.

### Private variants

We define single nucleotide variants (SNVs) as private (e.g. unique to individuals or families) in line with ACMG standards and guidelines [28] if the variant has not been recorded in any public database based on the Beacon Network (https://beacon-network.org/) after being corrected for batch effects. Such private SNVs were then additionally filtered to be coding and analysed with four orthogonal effect predictor methods CADD [29]), DANN [30], FATHMM-MKL [31] and ExAC-pLI [32] using default thresholds of 20, 0.95, 0.5 and 0.95, respectively to identify private SNVs with the highest possible confidence.

### Generation of reports

**The genome reports** were generated using variant calls derived from the WGS data as described above. A whole genome overview of the variant landscape of each participant was obtained by running the Variant Effect Predictor (VEP) v84 [33] with hg19 (GRCh37) cache. The called variants were interpreted in conjunction with public data from SNPedia [22], ExAC [32], GetEvidence [34] and ClinVar [35] for potentially beneficial and potentially harmful traits. A visual summary of the ancestry of each participant was obtained by merging the genotypes of each participant with genotypes from 2504 unrelated samples from 26 worldwide populations from the 1000 Genomes Project [36] and applying principal component analysis on the merged genotype matrix. Population membership proportions were inferred using the Admixture software [37] on the same genotype matrix.

**The methylome reports** were generated from the 450K data in conjunction with the epigenetic clock [38] for predictions on ageing and for predictions of exposure to smoking [39].

### Data access

All data reported here are available under open access from the PGP-UK data web page (https://www.personalgenomes.org.uk/data) which provides direct links to the corresponding public databases. However, as it is increasingly difficult to transfer data to the user, even under open access, there is a growing need for the analytics to be moved to where the relevant data are being stored. This concept is being addressed by cloud-based computing platforms e.g. through public-private partnerships offering a variety of models from open to fee-based data access [40-42], and easy access to training in big data analytics such as the online DataCamp programme [43]. Therefore, the reported PGP-UK data can also be accessed free of charge for non-commercial use on the Seven Bridges Cancer Genomics Cloud (CGC) [44], where PGP-UK data are hosted alongside relevant analyses tools enabling researchers to compute over these data in a cloud-based environment (http://www.cancergenomicscloud.org/).

### Genome app

GenoME was developed as an app for Apple iPads running iOS 9+. The app provides the public with a means to explore and better understand personal genomes. The app is fronted by four volunteer PGP-UK ambassadors, who share their personal genome stories through embedded videos and animated charts/information. All the features within the app that illustrate ancestry, traits and environmental exposures are populated by actual PGP-UK data from the corresponding participants. GenoME is freely available from the Apple App Store (https://itunes.apple.com/gb/app/genome/id1358680703?mt=8).

## Results

### Data types and access options

To demonstrate the feasibility of citizen science-driven contributions to personalized medicine, we actively engaged the first 10 participants and first three Genome Donors in all aspects of this PGP-UK pilot study. **Table 1** summarizes the matrix of 9 types of information (WGS, whole exome sequencing (WES), genotyping (e.g. 23andMe), 450K, WGBS, RNA-seq, Baseline Phenotypes, Reports and GenoME) which was generated for six categories (genome, methylome, transcriptome, phenotype, reports and GenoME app) and, where appropriate, the biological source from which the information was derived. The matrix comprises 103 datasets (∼2.5 TB) which were deposited according to data type in four different databases (ENA, EVA, ArrayExpress and PGP-UK), as there was no single public database able to host all data under open access. While easy access is facilitated through the PGP-UK data portal (see Links), the time required to download all the data can present a challenge that is common to many large-scale omics projects. The time to download all the datasets from **Table 1** using broadband with UK national average download speed of 36.2Mbps (according to official UK communication regulator Ofcom, 2017) would be more than 140 hours, indicating that faster solutions are required. To address this, we joined forces with Seven Bridges Genomics Inc (SBG), a leading provider of cloud-based computing infrastructure who pioneered such a platform for The Cancer Genome Atlas (TCGA). The Cancer Genomics Cloud (CGC, see Links) [44], funded as a pilot project by the US National Cancer Institute (NCI), allows academic researchers to access and collaborate on massive public cancer datasets, including the TCGA data. Researchers worldwide can create a free profile online or log in via their eRA Commons or NIH Center for Information Technology account to gain access to nearly 3 petabytes of publicly available data and relevant tools to analyse them. Following a successful trial and the open ethos of PGP, the data generated by the PGP-UK consortium for the first thirteen participants are now easily

**Table 1.**
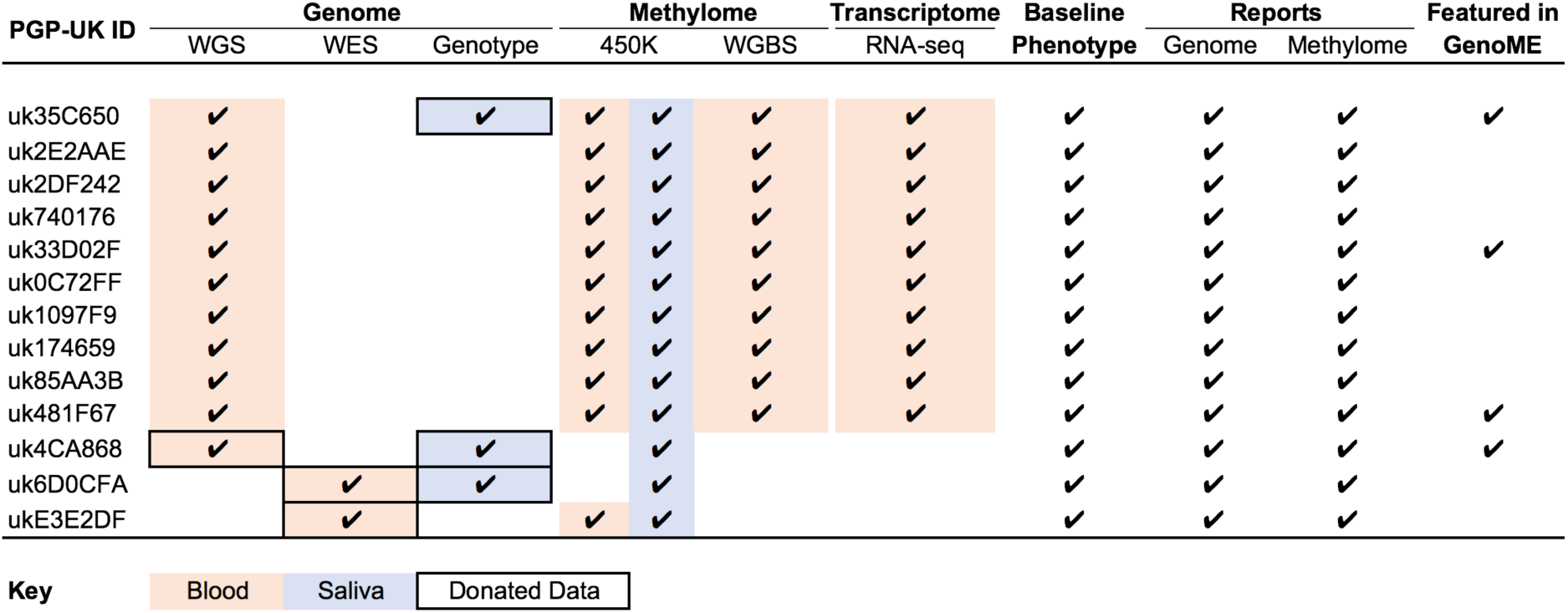
Information matrix of the PGP-UK pilot study. Ticks [✔] indicate the types of information available for each of the participants, the colour code depicts the biological source from which the information was derived and boxing highlights the information provided via Genome Donations. Genotype data are from 23andMe but other formats can also be donated.

accessible through the CGC for rapid, integrated and scalable cloud-based analysis using publicly available or custom-built pipelines.

### Genome reports

While data are the most useful information for the wider research community, reports were the most anticipated and intelligible information for the PGP-UK participants themselves. Great consideration was given to the content and format of the reports, taking on board valuable feedback from individual participants and the entire pilot group. At all times, the participants were made aware that both the data and reports were for information and research use only and not clinically accredited or actionable.

For the reporting of genetic variants, we opted for strict criteria so that the reported variants are low in number but as informative as possible (see Methods). On average, this resulted in over 200 incidental variants being reported for possibly beneficial and harmful traits. **Additional File 1** shows an exemplar genome report for participant uk35C650. In total, 4,105,373 SNVs were identified of which 97.5% were known and 2.5% (103,667 SNVs) appeared to be novel and thus private to this participant. Similar numbers were found for the other participants sequenced by PGP-UK, which is consistent with previous findings [36]. Of the known variants of participant uk35C650, 69 were associated with possibly beneficial traits (e.g. 6 SNVs associated with higher levels of high density lipoprotein (HDL) which is the ‘good’ type of cholesterol) and 193 with possibly harmful traits (e.g. 13 SNVs associated with Crohn’s disease, according to previously published studies). Taking advantage again of the open nature of PGP-UK, we shared the reports among all ten participants, which helped them to better understand the concept and meaning of beneficial or harmful SNV frequencies and distributions in the population. Since any genome report has the potential to uncover unexpected and even disconcerting information, the opportunity for participants to view other reports alongside their own provides context and reduces the likely anxiety if such reports are received and viewed in isolation. This was indeed confirmed as a positive aspect by all participants in the pilot study. In addition to learning about possibly beneficial and harmful variants, the participants were also interested to learn more about the ‘novel’ and potentially ‘private’ variants for which, by definition, nothing is yet known. This prompted us to investigate them in more detail.

### Private variants

A definition of what we consider private variants is described in the Methods section. **Table 2** shows the number of all, novel and private SNVs identified in ten of the participants using the PGP-UK analysis pipeline and additional, more stringent filtering against all openly accessible resources (see Methods). While this approach is imperfect due to some variants being represented in different ways and therefore easily missed [45], this effort reduced the number SNVs that are likely to be private to <20,000 per participant on average. To obtain a first insight into their possible functions we used multiple independent methods (see Methods and **Table 2**) to predict their effects. Of the 177,804 private SNVs identified, 29,558 (16.6%) passed the detection thresholds described in Methods and **Additional file 2**. As private SNVs cannot be validated in the traditional way, we used a Venn diagram (**Figure 1**) to assess the level of concordance/discordance between the four orthologous methods used. 47 SNVs were predicted to have significant impact by all four methods (**Figure 1**), providing the highest level of confidence that these are novel SNVs affecting gene function. Finally, we mapped these 47 private SNVs to their respective coding exons to reveal the affected genes (**Table 3**). The majority (41 SNVs) were predicted to have moderate impact, one was predicted to have high impact and four were predicted to have a modifier impact (**Table 3**).

**Figure 1.**
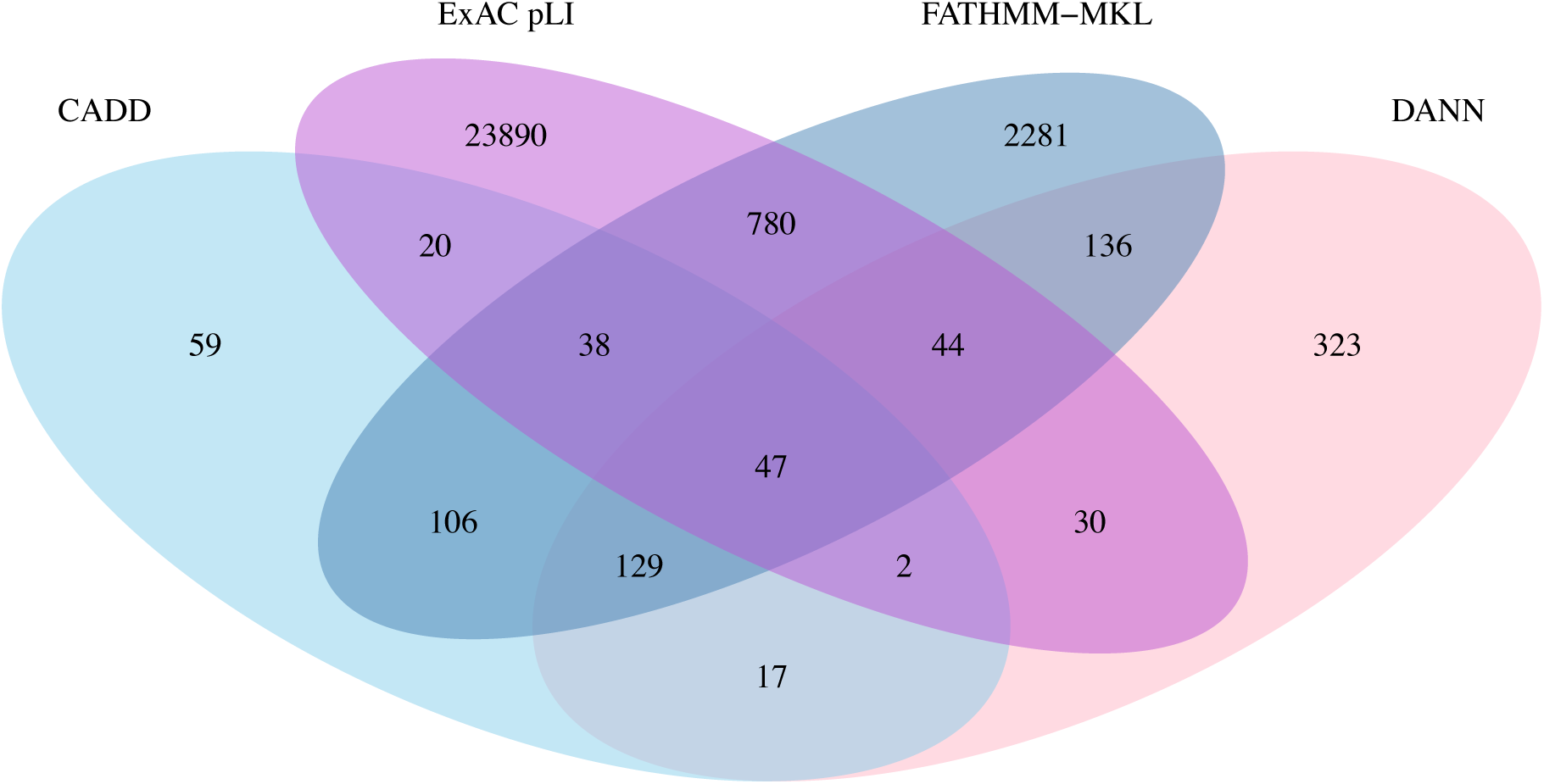
Venn diagram of private SNVs from the 10 PGP-UK pilot participants. Only coding SNVs were selected for effect prediction using the four orthogonal prediction methods indicated.

**Table 2.** Number of all, novel and private SNVs identified in the 10 PGP-UK pilot participants. Private SNVs were further analysed by four orthogonal methods for functional effects and the listed numbers reflect those SNVs that passed the thresholds described in Methods and **Additional file 2**.

|  | uk35C650 | uk2E2AAE | uk2DF242 | uk740176 | uk33D02F | uk0C72FF | uk1097F9 | uk174659 | uk85AA3B | uk481F67 | Total |
| --- | --- | --- | --- | --- | --- | --- | --- | --- | --- | --- | --- |
| **SNVs** |  |  |  |  |  |  |  |  |  |  |  |
| **All** | 4105831 | 4133496 | 4128205 | 4248904 | 4188095 | 4112224 | 4137176 | 4175392 | 4179265 | 4901623 | 42310211 |
| **Novel** | 103667 | 115090 | 131120 | 119803 | 119829 | 108299 | 112341 | 113893 | 118816 | 165833 | 1208691 |
| **Private** | 15020 | 14657 | 26420 | 22201 | 21208 | 13560 | 14093 | 14191 | 13983 | 22471 | 177804 |
| **Effect prediction** |  |  |  |  |  |  |  |  |  |  |  |
| **CADD** | 38 | 22 | 87 | 59 | 58 | 18 | 30 | 29 | 21 | 56 | 418 |
| **DANN** | 67 | 38 | 142 | 95 | 102 | 39 | 55 | 51 | 47 | 92 | 728 |
| **ExAC-pLI** | 2042 | 2113 | 3616 | 3115 | 2900 | 1909 | 2019 | 2069 | 1974 | 3094 | 24851 |
| **FATHMM-MKL** | 291 | 213 | 652 | 503 | 535 | 216 | 240 | 234 | 239 | 438 | 3561 |

**Table 3.** Genes predicted to be affected by private SNVs identified in the PGP-UK pilot.

| PGP-UK ID | SNV | Most Severe<br>Consequence Gene | Most Severe<br>Consequence | Impact |
| --- | --- | --- | --- | --- |
| uk481F67 | 1:235897198:C>A | LYST | Missense | Moderate |
| uk481F67 | 9:123540673:C>A | FBXW2 | Missense | Moderate |
| uk481F67 | 12:31595722:G>A | DENND5B | Missense | Moderate |
| uk35C650 | 4:126237966:G>T | FAT4 | Missense | Moderate |
| uk35C650 | 6:137019716:T>G | MAP3K5 | Missense | Moderate |
| uk35C650 | 11:18500391:G>T | LDHAL6A | Stop Gained | High |
| uk35C650 | 16:47494854:G>A | ITFG1 | Missense | Moderate |
| uk35C650 | 17:60767580:C>A | MRC2 | Missense | Moderate |
| uk35C650 | 20:34487400:G>A | PHF20 | Missense | Moderate |
| uk2E2AAE | 1:24687463:G>C | STPG1 | Missense | Moderate |
| uk2E2AAE | 22:40037056:G>T | CACNA1I | Missense | Moderate |
| uk2DF242 | 3:65342384:C>G | MAGI1 | Missense | Moderate |
| uk2DF242 | 3:180704791:T>C | DNAJC19 | Missense | Moderate |
| uk2DF242 | 4:48604097:C>G | FRYL | Missense | Moderate |
| uk2DF242 | 6:134638948:G>A | SGK1 | 5 Prime UTR | Modifier |
| uk2DF242 | 7:27867384:G>T | TAX1BP1 | Missense | Moderate |
| uk2DF242 | 9:27157892:A>T | TEK | Missense | Moderate |
| uk2DF242 | 9:32542215:T>C | TOPORS | Missense | Moderate |
| uk2DF242 | 12:108012140:G>A | BTBD11 | Missense | Moderate |
| uk2DF242 | 12:112617166:T>A | HECTD4 | Missense | Moderate |
| uk2DF242 | 14:103522914:G>A | CDC42BPB | Intron | Modifier |
| uk2DF242 | 16:30976974:G>C | SETD1A | Missense | Moderate |
| uk2DF242 | 17:29113033:G>T | CRLF3 | Missense | Moderate |
| uk2DF242 | 17:58503612:A>G | C17orf64 | Missense | Moderate |
| uk2DF242 | 19:6415701:C>T | KHSRP | Missense | Moderate |
| uk2DF242 | 19:34919334:C>T | UBA2 | 5 Prime UTR | Modifier |
| uk2DF242 | 20:37536816:G>C | PPP1R16B | Missense | Moderate |
| uk740176 | 4:114284614:A>C | ANK2 | Missense | Moderate |
| uk740176 | 7:130144866:A>T | MEST | Missense | Moderate |
| uk740176 | 10:94368844:C>A | KIF11 | Missense | Moderate |
| uk740176 | 19:8665875:C>G | ADAMTS10 | Missense | Moderate |
| uk33D02F | 16:69971109:T>G | WWP2 | Missense | Moderate |
| uk33D02F | 19:16640607:G>C | CHERP | Missense | Moderate |
| uk33D02F | X:49851274:C>T | CLCN5 | Missense | Moderate |
| uk0C72FF | 9:116034292:C>A | CDC26 | Intron | Modifier |
| uk1097F9 | 1:94640063:T>G | ARHGAP29 | Missense | Moderate |
| uk1097F9 | 7:105925660:C>T | NAMPT, RP11-22N19.2 | Upstream Gene | Modifier |
| uk1097F9 | 13:58207593:G>T | PCDH17 | Missense | Moderate |
| uk1097F9 | 17:30666952:G>A | C17orf75 | Missense | Moderate |
| uk1097F9 | 19:18234457:T>A | MAST3 | Missense | Moderate |
| uk1097F9 | 19:49655602:C>T | HRC | Missense | Moderate |
| uk174659 | 4:126238687:G>A | FAT4 | Missense | Moderate |
| uk174659 | 5:168114051:A>G | SLIT3 | Missense | Moderate |
| uk174659 | 10:93870886:C>T | CPEB3 | Missense | Moderate |
| uk85AA3B | 6:32945749:A>C | BRD2 | Missense | Moderate |
| uk85AA3B | 14:21840062:G>A | SUPT16H | Missense | Moderate |
| uk85AA3B | 15:52252874:C>T | LEO1 | Missense | Moderate |

### Methylome reports

There are currently no national or international policies or guidelines in place for the reporting of incidental epigenetic findings, including those based on DNA methylation [46], [47]. We limited our reports to categories for which findings had been independently validated and replicated, including the prediction of sex [38, 48], age [38] and smoking status [49]. **Additional file 3** shows an exemplar methylome report and **Table 4** summarizes our reported incidental epigenetic findings for the participants of the PGP-UK pilot. While the current methods for prediction of chronological age and sex are already well established and were highly accurate for all participants compared to the self-reported data, methods for an accurate prediction and interpretation of age deviation are still experimental. Averaged over two samples of different origin (blood and saliva), three of the thirteen participants showed significant age acceleration whereby the DNA methylation age is higher than the actual (chronological) age by more than 3.6 years, and three showed age deceleration (DNA methylation age is lower than the actual age by more than 3.6 years). Age deviation has already proved to be an informative biomarker. For instance, age acceleration has been reported to predict all-cause mortality in later life [50, 51] as well as cancer risk [52] and age deceleration has been linked to longevity [53, 54].

**Table 4.** Summary of 450K metthylome predictions for ten participants and three genome donations (boxed) of the PGP-UK pilot study. Predictions were made for sex, age and smoking status using blood or saliva as indicated. Asterisks (*) denote that methylation age acceleration and deceleration is only present if methylation age is more than 3.6 years different to the respective actual age [50].

| PGP-UK ID | uk35C650 | uk2E2AAE | uk2DF242 | uk740176 | uk33D02F | uk0C72FF | uk1097F9 | uk174659 | uk85AA3B | uk481F67 | uk4CA868 | uk6D0CFA | ukE3E2DF |
| --- | --- | --- | --- | --- | --- | --- | --- | --- | --- | --- | --- | --- | --- |
| **Sex prediction** |  |  |  |  |  |  |  |  |  |  |  |  |  |
| **Actual sex** | Male | Female | Male | Female | Female | Male | Male | Male | Female | Male | Male | Male | Male |
| **Predicted sex** | Male | Female | Male | Female | Female | Male | Male | Male | Female | Male | Male | Male | Male |
| **Age prediction** |  |  |  |  |  |  |  |  |  |  |  |  |  |
| **Actual age at sampling** | 59.67 | 56.49 | 48.93 | 47.94 | 44.57 | 64.7 | 50.97 | 33.65 | 50.59 | 57.45 | 57.03 | 39.77 | 32.83 |
| **Estimated methylation age (blood)** | 52.78 | 53.21 | 47.52 | 52.61 | 43.45 | 56.19 | 45.23 | 41.7 | 54.32 | 63.07 | NA | NA | 43.67 |
| **Estimated methylation age (saliva)** | 53.49 | 52.22 | 47.93 | 53.3 | 42.15 | 61.28 | 50.74 | 35.53 | 53.29 | 58.47 | 56.18 | 37.85 | 37.98 |
| **Average methylation age** | 53.14 | 52.72 | 47.73 | 52.96 | 42.8 | 58.74 | 47.99 | 38.62 | 53.81 | 60.77 | NA | NA | 40.83 |
| **Methylation age acceleration\*** | None | None | None | 5.02 years | None | None | None | 4.97 years | None | None | None | None | 8.00 years |
| **Methylation age deceleration\*** | 6.53 years | 3.77 years | None | None | None | 5.96 years | None | None | None | None | None | None | None |
| **Smoking status prediction** |  |  |  |  |  |  |  |  |  |  |  |  |  |
| **Actual smoking status** | Past | Never | Never | Never | Never | Occasional | Never | Never | Never | Past | Past | Past | Never |
| **Smoking score (saliva)** | 1.55 | 1.32 | -3.21 | -3.28 | -1.78 | 0.67 | 3.35 | 0.25 | -1.84 | 7.04 | 3.86 | -2.54 | 2.8 |
| **smoking score (blood)** | -1.6 | -2 | -5.63 | -4.32 | -2.7 | -2.4 | -2.08 | -3.53 | -4.68 | 1.26 | NA | NA | -4.02 |
| **smoking prediction** | Past/ Never | Past/ Never | Past/ Never | Past/ Never | Past/ Never | Past/ Never | Past/ Never | Past/ Never | Past/ Never | Past/ Never | Past/Never | Past/Never | Past/Never |

The final category which was reported back to participants was exposure to smoking. Epigenetic associations with environmentally mediated exposures are typically measured through epigenome-wide association studies (EWAS) [55]. Based on the analysis of DNA methylation in saliva and blood samples, smoking scores were generated using the method developed by Elliott et al. [39]. The smoking score was calculated using weighted methylation values of 187 well established smoking-associated CpG sites [56] and has been shown to accurately predict whether individuals are past/never or current smokers [39]. This study showed that a smoking score of more than 17.55 for Europeans, or more than 11.79 for South Asians indicated that an individual is a current smoker, while values below these thresholds indicate that individuals are past or never smokers. All participants in the PGP-UK pilot study were predicted to be past or never smokers, in both saliva and blood samples. The prediction was correct for 12 out of the 13 participants who self-reported as either past or never smokers. However, one participant (uk0C72FF) self-reported as an ‘occasional smoker’. This aberrant prediction could be explained by the study population in which the threshold was set; the individuals considered ‘current smokers’ smoked a mean of 23 cigarettes per day for Europeans and 13 per day for South Asians. Consequently, very occasional smoking may not classify as ‘current smoking’ using the thresholds of Elliott et al. [39]. Another limitation is that the smoking score was tested in European and South Asian populations, thus it may be less accurate in other ethnicities.

### GenoME app

To make genome and methylome reports more accessible and understandable to the lay public, we developed GenoME as a free and open source genome app for Apple iPads. The main purpose was to have actual people presenting real incidental findings in an innovative and engaging way. For that, we recruited four volunteers (ambassadors) from the pilot cohort who were willing to self-identify and share their personal genome story through embedded videos, specifically composed music and artistically animated examples of incidental findings from their genomes. To illustrate this, we selected two traits (eye colour and smoking status) for which we reported genetic and epigenetic variants, respectively. **Figure 2** shows three screen shots of how SNVs associated with eye colour are communicated. **Figure 2A** shows one of the ambassadors and explanatory text in the left panel and a whirling cloud of colour representing all possible eye colours in the right panel. **Figure 2B** shows an intermediate state of the colours coalescing into the eye colour predicted by the SNVs for this participant and **Figure 2C** shows the final stage of zooming in on the ambassador’s actual eye colour for comparison with the predicted eye colour which was correct in this case. In GenoME, the sequence of screens is complemented by integrated music elements to enable people with compromised sight to experience genetic variation through sound. **Figure 3** shows a similar sequence of three screens for the prediction smoking status based on epigenetic (DNA methylation) variants. In this case, a cloud of smoke coalesces into ‘never/past’ or ‘current’ smoker icons depending on the epigenetic profile of the participant. Other features (not shown) include variants associated with ancestry using an animated world map and disease using population-specific allele frequency graphics.

**Figure 2.**
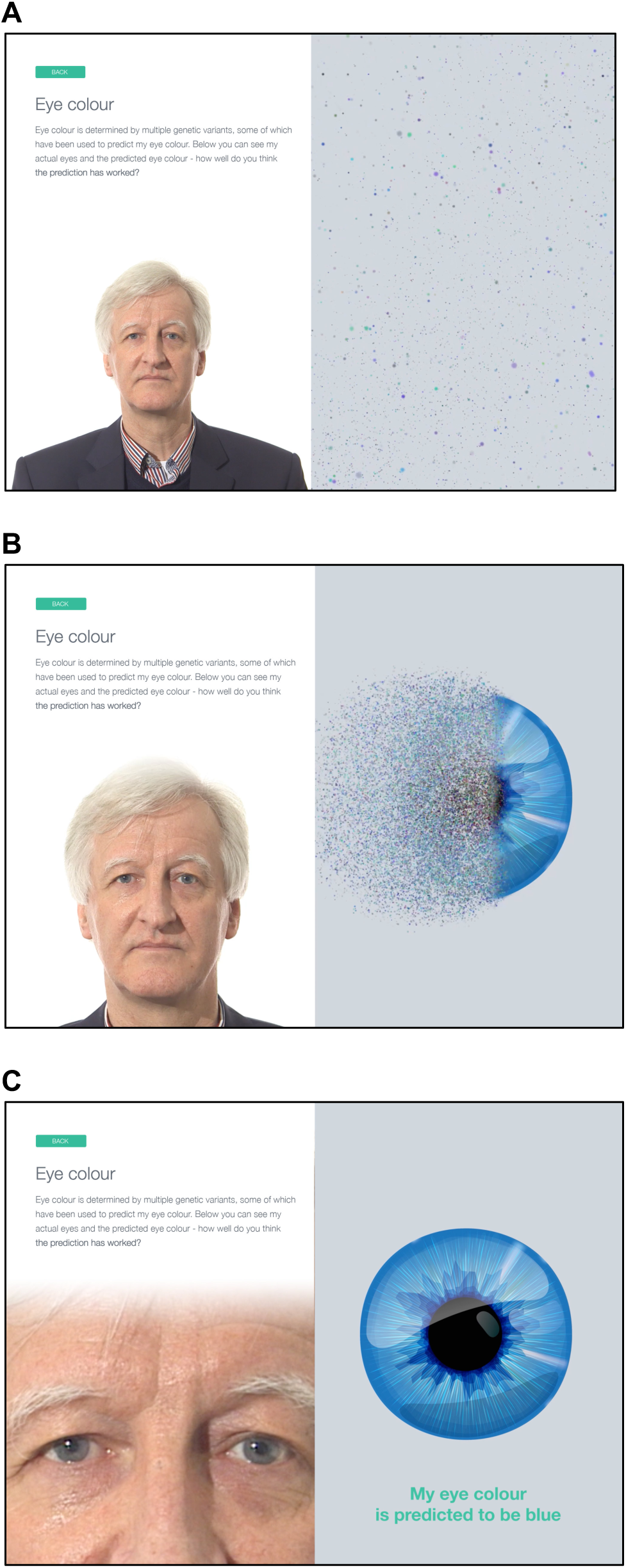
Time series (A-C) of screens showing how GenoME communicates genetic SNVs associated with the participant’s eye colour.

**Figure 3.**
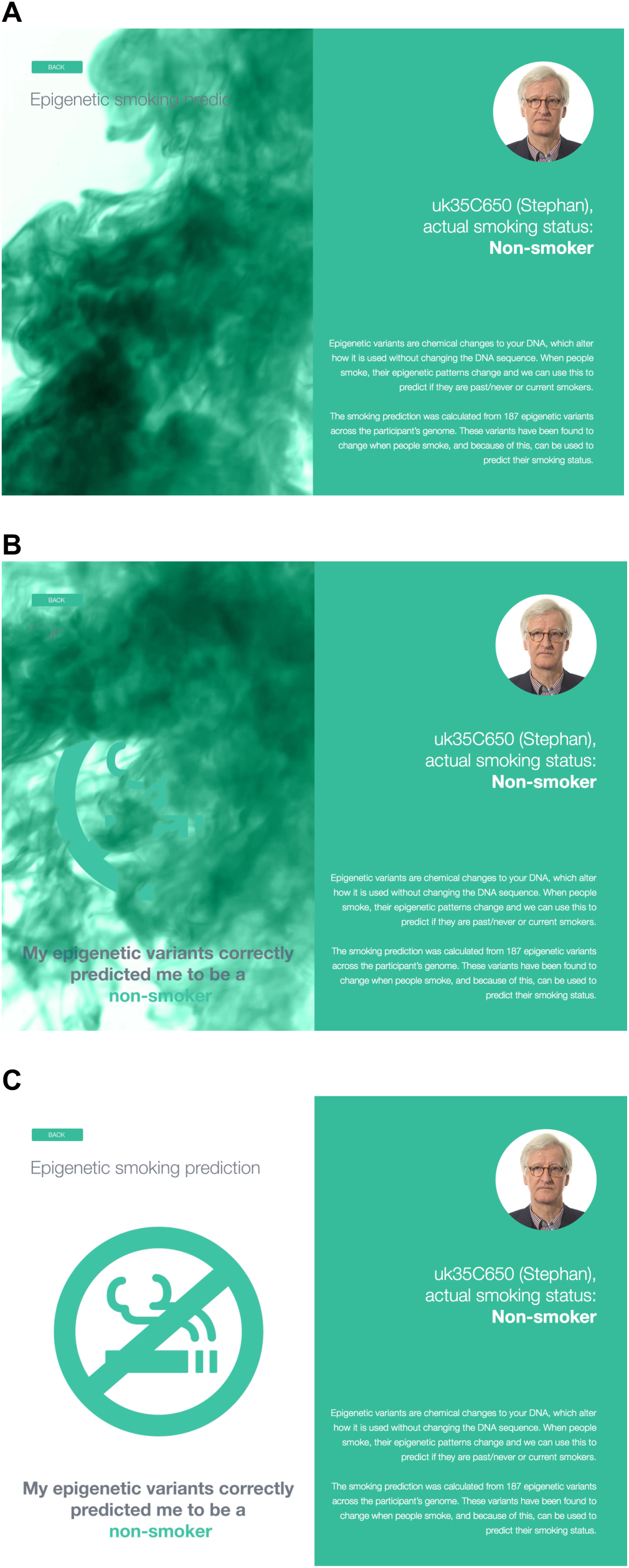
Time series (A-C) of screens showing how GenoME communicates epigenetic SNVs associated with the participant’s smoking status.

## Discussion

In this study, we report the study design, data processing and findings of the PGP-UK pilot, and demonstrate the suitability of PGP-UK as a hybrid between a research and a citizen science project. For the latter, we enlisted 11 citizen scientists who made up a third of the named authors and contributed vitally to the assessment of our reporting strategy, features of GenoME and advocacy of citizen science in general. As part of our citizen science programme, PGP-UK encourages such interactions also on the international level through membership of the global PGP Network, DNA.Land [57] and Open Humans (see Links), a project which enables individuals to connect their data with research and citizen science worldwide.

The resource value of PGP-UK will become more apparent as more participants are enrolled and data released. Towards this, a second batch of data and reports has already been released (see Links) for another 94 participants and the ultimate goal is to eventually reach the 100K participants mark for which ethics approval has been obtained. Considering the scale of past and on-going UK sequencing projects [4, 58], we believe this is achievable especially though utilizing the genome donation procedure described here. In the meantime, the PGP-UK data also contribute to the global PGP resource. According to Repositive (see Links), a platform linking open access data across 49 resources, the global PGP network has collectively generated over 1,121 data sets. Additionally and in the N=1 context of personalised medicine, each data set is of course highly informative in its own right [59].

To our knowledge, the methylome reports described here are the first of their kind issued for any incidental epigenetic findings. The open nature of PGP-UK makes it possible to explore appropriate frameworks and guidelines in a controlled environment [46, 47]. Based on our experience, there is high interest and acceptance for adequately validated and replicated epigenetic findings to be reported back alongside genetic findings, particularly those that capture environmental exposures such as tobacco smoke, alcohol consumption and air pollution. Accordingly, we are now evaluating if any EWAS-derived variants are yet appropriate for inclusion. Another area of potential future interest is the prediction of a participant’s suitability as donor in the context of transplant medicine [60]. Another innovation reported here is GenoME, an app for exploring personal genomes. Apps can easily reach millions of people and thus are an ideal stepping-stone to engage with citizen science, which plays an important role in making personal and medical genomics acceptable to the public. A recent study – Your DNA, Your Say – concluded that “*Genomic medicine can only be successfully integrated into healthcare if there are aligned programmes of engagement that enlighten the public to what genomics is and what it can offer*” [61]. This is particularly important as we reach the cusp of widespread implementation of genomic and personalised medicine.

Our study also highlighted some limitations. For instance, adequate public databases for multi-dimensional open access data, equivalent to dbGaP [62] or EGA [63] for controlled access data, are currently still lacking. Consequently, the PGP-UK data were submitted to multiple open access public databases, depending on type of data. Furthermore, most public databases are not built to host or enable downloading of TB-scale datasets and don’t enable easy access and analysis without downloading the data. To overcome these current limitations, we made the PGP-UK pilot data also available in a cloud-based system [44]. The high level of automation implemented for the PGP-UK analysis pipeline allows updates to be generated and released as and when required and new reports (e.g. based on WGBS and RNA-seq data) to be added in the future. At the time of submission, over 100 genomes and associated reports had been generated, released (see Links) and deposited into public databases.

## Conclusion

Our results demonstrate that omics-based research and citizen science can be successfully hybridised into an open access resource for personal and medical genomics. The key features that allowed this were transparency and interoperability on the people and data levels, resulting in a degree of openness that is not generally found in medical research and thus provides an alternative to traditional research models. The introduction of the GenoME app and the framework for Genome Donations provide two novel modes for the public to engage with personal and medical genomics.

## Acknowledgments

PGP-UK gratefully acknowledges voluntary contributions from the researchers and participants, technical assistance from Harvard PGP Team, the UCL/UCLH Biobank for Studying Health and Disease for DNA/RNA preparation and UCL Genomics for array processing and funding support from the UCL Cancer Institute Research Trust, the Frances and Augustus Newman Foundation, Dangoor Education and the National Institute for Health Research UCLH Biomedical Research Centre (BRC369/CN/SB/101310). The development of the GenoME app was supported by a donation from Michael Chowen CBE DL and Maureen Chowen. We also would like to acknowledge an award from the MRC Proximity to Discovery Industry Engagement Fund (530912) to facilitate access to the cloud-based computing infrastructure of Seven Bridges Genomics.

## Project coordination

Stephan Beck, Olga Chervova, Lucia Conde, José Afonso Guerra-Assunção, Rifat Hamoudi, Paul L Harrison, Javier Herrero, Erica Jones, Jane Kaye, Ismail Moghul, Amy P Webster

## IT systems

Olga Chervova, Ricarda Gaentzsch, Javier Herrero, Rifat Hamoudi, Vincent Harding, Ismail Moghul, Sevgi Umur

## Data Analysis

Lucia Conde, Simone Ecker, Silvana A Fioramonti, José Afonso Guerra-Assunção, Rifat Hamoudi, Javier Herrero, Emanuele Libertini, Ismail Moghul, Nikolas Pontikos, Kirti Rana, Amy P Webster

## Citizen science

Graham Bignell, Maggie Bond, Martin J Callanan, Manuel Corpas, Deirdre Gribbin, Laura McCormack, Nikolas Pontikos, Momodou Semega-Janneh, Colin P Smith, Karen Wint, John N Wood

## Manuscript

Stephan Beck wrote the manuscript with contributions from all authors. All authors have approved the manuscript. Corresponding author: Stephan Beck.

## Links

PGP-UK: https://www.personalgenomes.org.uk/

PGP-UK Data: https://www.personalgenomes.org.uk/data/

GenoME app: https://itunes.apple.com/gb/app/genome/id1358680703?mt=8

Global PGP Network: https://www.personalgenomes.org.uk/global-network

Open Humans: https://www.openhumans.org/

DNA.Land https://dna.land/

EBI: http://www.ebi.ac.uk/

Repositive: https://discover.repositive.io/collections/

Seven Bridges Genomics: https://www.sevenbridges.com/

Cancer Genomics Cloud (access to PGP-UK and other publicly available data): http://www.cancergenomicscloud.org/

1000 Genomes Project: www.internationalgenome.org/

SNPedia: www.snpedia.com/

ExAC: http://exac.broadinstitute.org/

GetEvidence: http://evidence.pgp-hms.org/

ClinVar: www.ncbi.nlm.nih.gov/clinvar/

## References

1. Lander ES, Linton LM, Birren B, Nusbaum C, Zody MC, Baldwin J, Devon K, Dewar K, Doyle M, FitzHugh W, et al: **Initial sequencing and analysis of the human genome**. Nature 2001, 409:860–921.

2. Venter JC, Adams MD, Myers EW, Li PW, Mural RJ, Sutton GG, Smith HO, Yandell M, Evans CA, Holt RA, et al: **The sequence of the human genome**. Science 2001, 291:1304–1351.

3. Stephens ZD, Lee SY, Faghri F, Campbell RH, Zhai C, Efron MJ, Iyer R, Schatz MC, Sinha S, Robinson GE: **Big Data: Astronomical or Genomical?** PLoS Biol 2015, 13:e1002195.

4. Peplow M: The 100,000 Genomes Project. BMJ 2016, 353:i1757.

5. Collins FS, Varmus H: **A new initiative on precision medicine**. N Engl J Med 2015, 372:793–795.

6. Su P: Direct-to-consumer genetic testing: a comprehensive view. Yale J Biol Med 2013, 86:359–365.

7. Greenbaum D, Sboner A, Mu XJ, Gerstein M: Genomics and privacy: implications of the new reality of closed data for the field. PLoS Comput Biol 2011, 7:e1002278.

8. Reardon J, Ankeny RA, Bangham J, K WD, Hilgartner S, Jones KM, Shapiro B, Stevens H, Genomic Open workshop g: **Bermuda 2.0: reflections from Santa Cruz**. Gigascience 2016, 5:1–4.

9. Ball MP, Bobe JR, Chou MF, Clegg T, Estep PW, Lunshof JE, Vandewege W, Zaranek A, Church GM: **Harvard Personal Genome Project: lessons from participatory public research**. Genome Med 2014, 6:10.

10. Ball MP, Thakuria JV, Zaranek AW, Clegg T, Rosenbaum AM, Wu XD, Angrist M, Bhak J, Bobe J, Callow MJ, et al: **A public resource facilitating clinical use of genomes**. Proceedings of the National Academy of Sciences of the United States of America 2012, 109:11920–11927.

11. Mao Q, Ciotlos S, Zhang RY, Ball MP, Chin R, Carnevali P, Barua N, Nguyen S, Agarwal MR, Clegg T, et al: **The whole genome sequences and experimentally phased haplotypes of over 100 personal genomes**. Gigascience 2016, 5:42.

12. Reuter MS, Walker S, Thiruvahindrapuram B, Whitney J, Cohn I, Sondheimer N, Yuen RKC, Trost B, Paton TA, Pereira SL, et al: **The Personal Genome Project Canada: findings from whole genome sequences of the inaugural 56 participants**. CMAJ 2018, 190:E126–E136.

13. Chen R, Mias GI, Li-Pook-Than J, Jiang L, Lam HY, Chen R, Miriami E, Karczewski KJ, Hariharan M, Dewey FE, et al: **Personal omics profiling reveals dynamic molecular and medical phenotypes**. Cell 2012, 148:1293–1307.

14. Becnel LB, Pereira S, Drummond JA, Gingras MC, Covington KR, Kovar CL, Doddapaneni HV, Hu J, Muzny D, McGuire AL, et al: **An open access pilot freely sharing cancer genomic data from participants in Texas**. Sci Data 2016, 3:160010.

15. Dyke SO, Kirby E, Shabani M, Thorogood A, Kato K, Knoppers BM: **Registered access: a ’Triple-A’ approach**. Eur J Hum Genet 2016, 24:1676–1680.

16. Lunshof JE, Chadwick R, Vorhaus DB, Church GM: **From genetic privacy to open consent**. Nat Rev Genet 2008, 9:406–411.

17. Woolley JP, McGowan ML, Teare HJ, Coathup V, Fishman JR, Jr RAS, Sterckx S, Kaye J, Juengst ET: Citizen science or scientific citizenship? Disentangling the uses of public engagement rhetoric in national research initiatives. BMC Medical Ethics 2016.

18. Boulos MN, Brewer AC, Karimkhani C, Buller DB, Dellavalle RP: **Mobile medical and health apps: state of the art, concerns, regulatory control and certification**. Online J Public Health Inform 2014, 5:229.

19. Vassy JL, Christensen KD, Schonman EF, Blout CL, Robinson JO, Krier JB, Diamond PM, Lebo M, Machini K, Azzariti DR, et al: **The Impact of Whole-Genome Sequencing on the Primary Care and Outcomes of Healthy Adult Patients: A Pilot Randomized Trial**. Ann Intern Med 2017.

20. Li H, Durbin R: Fast and accurate short read alignment with Burrows-Wheeler transform. Bioinformatics 2009, 25:1754–1760.

21. Li H, Handsaker B, Wysoker A, Fennell T, Ruan J, Homer N, Marth G, Abecasis G, Durbin R, Genome Project Data Processing S: **The Sequence Alignment/Map format and SAMtools**. Bioinformatics 2009, 25:2078–2079.

22. Cariaso M, Lennon G: SNPedia: a wiki supporting personal genome annotation, interpretation and analysis. Nucleic Acids Res 2012, 40:D1308–1312.

23. Merkel A, Fernandez-Callejo M, Casals E, Marco-Sola S, Schuyler R, Gut IG, Heath SC: GEMBS — high through-put processing for DNA methylation data from Whole Genome Bisulfite Sequencing (WGBS). bioRxiv 2017.

24. Suzuki M, Liao M, Wos F, Johnston AD, DeGrazia J, Ishii J, Bloom T, Zody MC, Germer S, Greally JM: Whole genome bisulfite sequencing using the Illumina HiSeq X system. BioRxiv 2017.

25. Morris TJ, Butcher LM, Feber A, Teschendorff AE, Chakravarthy AR, Wojdacz TK, Beck S: **ChAMP: 450k Chip Analysis Methylation Pipeline**. Bioinformatics 2014, 30:428–430.

26. Yuan Tian TJM, Amy P Webster, Zhen Yang, Stephan Beck, Andrew Feber, Andrew E Teschendorff: **ChAMP: Updated Methylation Analysis Pipeline for Illumina BeadChips**. Bioinformatics 2017.

27. Aryee MJ, Jaffe AE, Corrada-Bravo H, Ladd-Acosta C, Feinberg AP, Hansen KD, Irizarry RA: **Minfi: a flexible and comprehensive Bioconductor package for the analysis of Infinium DNA methylation microarrays**. Bioinformatics 2014, 30:1363–1369.

28. Richards S, Aziz N, Bale S, Bick D, Das S, Gastier-Foster J, Grody WW, Hegde M, Lyon E, Spector E, et al: Standards and guidelines for the interpretation of sequence variants: a joint consensus recommendation of the American College of Medical Genetics and Genomics and the Association for Molecular Pathology. Genet Med 2015, 17:405–424.

29. Kircher M, Witten DM, Jain P, O’Roak BJ, Cooper GM, Shendure J: **A general framework for estimating the relative pathogenicity of human genetic variants**. Nat Genet 2014, 46:310–315.

30. Quang D, Chen Y, Xie X: DANN: a deep learning approach for annotating the pathogenicity of genetic variants. Bioinformatics 2015, 31:761–763.

31. Shihab HA, Gough J, Cooper DN, Stenson PD, Barker GL, Edwards KJ, Day IN, Gaunt TR: Predicting the functional, molecular, and phenotypic consequences of amino acid substitutions using hidden Markov models. Hum Mutat 2013, 34:57–65.

32. Lek M, Karczewski KJ, Minikel EV, Samocha KE, Banks E, Fennell T, O’Donnell-Luria AH, Ware JS, Hill AJ, Cummings BB, et al: **Analysis of protein-coding genetic variation in 60,706 humans**. Nature 2016, 536:285–291.

33. McLaren W, Gil L, Hunt SE, Riat HS, Ritchie GR, Thormann A, Flicek P, Cunningham F: **The Ensembl Variant Effect Predictor**. Genome Biol 2016, 17:122.

34. Ball MP, Thakuria JV, Zaranek AW, Clegg T, Rosenbaum AM, Wu X, Angrist M, Bhak J, Bobe J, Callow MJ, et al: **A public resource facilitating clinical use of genomes**. Proc Natl Acad Sci U S A 2012, 109:11920–11927.

35. Landrum MJ, Lee JM, Benson M, Brown G, Chao C, Chitipiralla S, Gu B, Hart J, Hoffman D, Hoover J, et al: **ClinVar: public archive of interpretations of clinically relevant variants**. Nucleic Acids Res 2016, 44:D862–868.

36. Genomes Project C, Auton A, Brooks LD, Durbin RM, Garrison EP, Kang HM, Korbel JO, Marchini JL, McCarthy S, McVean GA, Abecasis GR: **A global reference for human genetic variation**. Nature 2015, 526:68–74.

37. Alexander DH, Novembre J, Lange K: Fast model-based estimation of ancestry in unrelated individuals. Genome Res 2009, 19:1655–1664.

38. Horvath S: DNA methylation age of human tissues and cell types. Genome Biol 2013, 14:R115.

39. Elliott HR, Tillin T, McArdle WL, Ho K, Duggirala A, Frayling TM, Davey Smith G, Hughes AD, Chaturvedi N, Relton CL: **Differences in smoking associated DNA methylation patterns in South Asians and Europeans**. Clin Epigenetics 2014, 6:4.

40. Molnar-Gabor F, Lueck R, Yakneen S, Korbel JO: Computing patient data in the cloud: practical and legal considerations for genetics and genomics research in Europe and internationally. Genome Med 2017, 9:58.

41. Granados Moreno P, Joly Y, Knoppers BM: Public-Private Partnerships in Cloud-Computing Services in the Context of Genomic Research. Front Med (Lausanne) 2017, 4:3.

42. Langmead B, Nellore A: **Cloud computing for genomic data analysis and collaboration**. Nature Review Genetics 2018, 19:208–220.

43. Munevar S: Unlocking Big Data for better health. Nat Biotechnol 2017, 35:684–686.

44. Lau JW, Lehnert E, Sethi A, Malhotra R, Kaushik G, Onder Z, Groves-Kirkby N, Mihajlovic A, DiGiovanna J, Srdic M, et al: **The Cancer Genomics Cloud: Collaborative, Reproducible, and Democratized-A New Paradigm in Large-Scale Computational Research**. Cancer Res 2017, 77:e3–e6.

45. Bayat A, Gaeta B, Ignjatovic A, Parameswaran S: **Improved VCF normalization for accurate VCF comparison**. Bioinformatics 2017, 33:964–970.

46. Carter AC, Chang HY, Church G, Dombkowski A, Ecker JR, Gil E, Giresi PG, Greely H, Greenleaf WJ, Hacohen N, et al: **Challenges and recommendations for epigenomics in precision health**. Nat Biotechnol 2017, 35:1128–1132.

47. Dyke SOM, Saulnier KM, Dupras C, Procaccini D, Webster AP, Maschke K, Rothstein M, Siebert R, Walter J, Beck S, et al: Points-to-Consider on the Return of Results in Epigenetic Research. under revision.

48. Liu J, Morgan M, Hutchison K, Calhoun VD: **A study of the influence of sex on genome wide methylation**. PLoS One 2010, 5:e10028.

49. Gao X, Jia M, Zhang Y, Breitling LP, Brenner H: DNA methylation changes of whole blood cells in response to active smoking exposure in adults: a systematic review of DNA methylation studies. Clin Epigenetics 2015, 7:113.

50. Marioni RE, Shah S, McRae AF, Chen BH, Colicino E, Harris SE, Gibson J, Henders AK, Redmond P, Cox SR, et al: **DNA methylation age of blood predicts all-cause mortality in later life**. Genome Biol 2015, 16:25.

51. Chen BH, Marioni RE, Colicino E, Peters MJ, Ward-Caviness CK, Tsai PC, Roetker NS, Just AC, Demerath EW, Guan W, et al: **DNA methylation-based measures of biological age: meta-analysis predicting time to death**. Aging (Albany NY) 2016, 8:1844–1865.

52. Yang Z, Wong A, Kuh D, Paul DS, Rakyan VK, Leslie RD, Zheng SC, Widschwendter M, Beck S, Teschendorff AE: **Correlation of an epigenetic mitotic clock with cancer risk**. Genome Biol 2016, 17:205.

53. McEwen LM, Morin AM, Edgar RD, MacIsaac JL, Jones MJ, Dow WH, Rosero-Bixby L, Kobor MS, Rehkopf DH: **Differential DNA methylation and lymphocyte proportions in a Costa Rican high longevity region**. Epigenetics Chromatin 2017, 10:21.

54. Horvath S, Gurven M, Levine ME, Trumble BC, Kaplan H, Allayee H, Ritz BR, Chen B, Lu AT, Rickabaugh TM, et al: **An epigenetic clock analysis of race/ethnicity, sex, and coronary heart disease**. Genome Biol 2016, 17:171.

55. Rakyan VK, Down TA, Balding DJ, Beck S: **Epigenome-wide association studies for common human diseases**. Nat Rev Genet 2011, 12:529–541.

56. Zeilinger S, Kuhnel B, Klopp N, Baurecht H, Kleinschmidt A, Gieger C, Weidinger S, Lattka E, Adamski J, Peters A, et al: **Tobacco smoking leads to extensive genome-wide changes in DNA methylation**. PLoS One 2013, 8:e63812.

57. Yuan J, Gordon A, Speyer D, Aufrichtig R, Zielinski D, Pickrell J, Erlich Y: **DNA.****Land is a framework to collect genomes and phenomes in the era of abundant genetic information**. Nat Genet 2018, 50:160–165.

58. Kaye J, Hurles M, Griffin H, Grewal J, Bobrow M, Timpson N, Smee C, Bolton P, Durbin R, Dyke S, et al: **Managing clinically significant findings in research: the UK10K example**. Eur J Hum Genet 2014, 22:1100–1104.

59. Lillie EO, Patay B, Diamant J, Issell B, Topol EJ, Schork NJ: **The n-of-1 clinical trial: the ultimate strategy for individualizing medicine?** Per Med 2011, 8:161–173.

60. Paul DS, Jones A, Sellar RS, Mayor NP, Feber A, Webster AP, Afonso N, Sergeant R, Szydlo RM, Apperley JF, et al: A donor-specific epigenetic classifier for acute graft-versus-host disease severity in hematopoietic stem cell transplantation. Genome Med 2015, 7:128.

61. Middleton A: **Your DNA, Your Say**. New Bioeth 2017, 23:74–80.

62. Tryka KA, Hao L, Sturcke A, Jin Y, Wang ZY, Ziyabari L, Lee M, Popova N, Sharopova N, Kimura M, Feolo M: **NCBI’s Database of Genotypes and Phenotypes: dbGaP**. Nucleic Acids Res 2014, 42:D975–979.

63. Hoogstrate Y, Zhang C, Senf A, Bijlard J, Hiltemann S, van Enckevort D, Repo S, Heringa J, Jenster G, R JAF, et al: **Integration of EGA secure data access into Galaxy**. F1000Res 2016, 5.

