## Supplementary Materials for "PGP-UK: a research and citizen science hybrid project in support of personalized medicine"

*A list of authors, their affiliations and contributions appears at the end of the paper.

#### Supplementary Material

### Additional Files

**Additional file 1.** Exemplar PGP-UK genome report of participant uk35C650 providing details on the number and type of variants and their association with ancestry as well as possibly beneficial and harmful traits. The released versions of the reports provide links to the underlying databases, including SNPedia, ExAC, GetEvidence and ClinVar.

#### Genomics Report for PGP-UK1/uk35C650

##### 1 Summary

This is the genome report for participant PGP-UK1/uk35C650 . It was produced using collaborative research tools, including SNPedia and GetEvidence. This summary shows an overview of all the variants which were found in the genome for this individual. They have been compared with a reference genome.

This report was generated automatically and is not clinically approved. It is provided for personal and research purposes only.

This document contains hyperlinks, shown in grey, that will take you to external websites where you can find more detailed explanations. Some of the technical terms are also explained in more detail in the Ensembl Glossary. We would welcome your feedback about this report, for example, if you would like more information about anything or if any of the links have become inactive. You can contact us on:.

This summary shows an overview of all the variants which were found in the genome for this individual. The "variants remaining after filtering" refers to any differences in the DNA identified when compared to the reference genome. Of these, the majority will have already been found in some other sequenced individual and put on a database (existing variants) while others have not yet been annotated (novel variants).

"Overlapped genes" refers to the number of times where a variant was found in a region of the genome containing a gene. "Exon" refers to the part of the gene which goes on to form a protein, and variants in this part of the gene are more likely to cause changes in the shape of the protein. Upstream, downstream, intronic and intergenic variants are more likely to alter the regulation of that gene but will not change the protein itself.

A transcript for a protein-coding gene can include the exons, introns and other gene features that are transcribed and important for gene function but might not be translated into the final protein. Not all transcripts are for protein-coding genes, with many containing non-coding RNAs that can be overlapping other genes, in introns or in intergenic regions. The diagram in Figure 1 is a simplification of the usual gene structure.

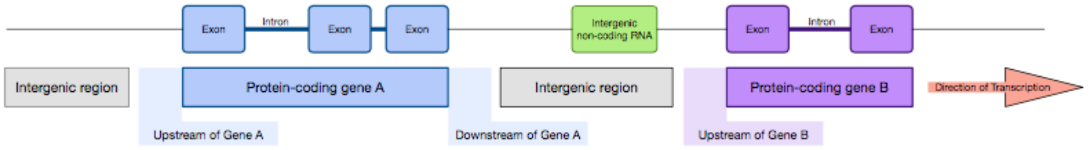


Figure 1: Diagram of gene structure indicating locations of potential variants

| Feature | Count |
|---|---|
| Lines of input read | 4130989 |
| Variants remaining after filtering | 4105273 |
| Novel / existing variants | 103667 (2.5%) / 4001606 (97.5%) |
| Overlapped genes | 54674 |
| Overlapped transcripts | 64349 |
| Overlapped regulatory features | 211555 |

Table 1: Variant calling summary

There are several different types of genomic variants. The most common are single nucleotide variants (SNV) that correspond to the change of a single nucleotide in the DNA. Other variant types include insertions, where the DNA in the individual is longer than the reference sequence due to the insertion of one or more nucleotides; and deletions, where a few nucleotides are missing compared to the reference sequence.

Some of these changes will have no effect on the protein, while some changes may alter the protein function to varying degrees. The PolyPhen analysis software attempts to quantify the effect each mutation will have on the protein function. This ranges from "benign" where no change to the protein function is expected, to "probably damaging" where it is predicted that the mutation will affect protein function. It is nevertheless important to note that what is "damaging" for the protein is not necessarily damaging for the individual.

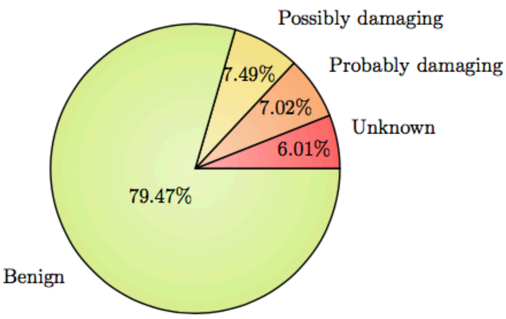


Figure 2: PolyPhen Summary

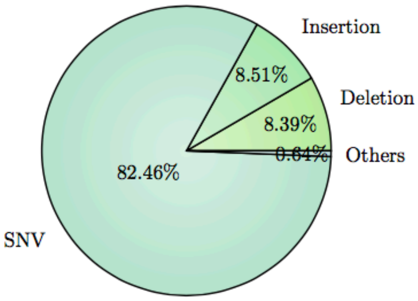


Figure 3: Variant Class

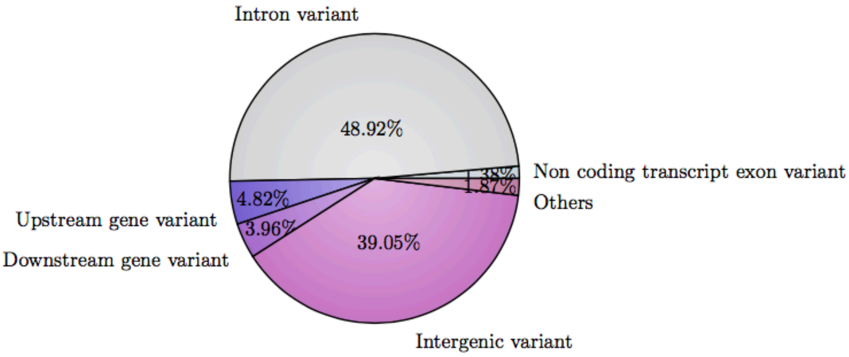


Figure 4: Consequence type

#### 2 Ancestry

This plot shows the distribution of the genomes of different populations. Data from several studies which used whole genome sequencing was used to see the relationships between the genomes of the populations. It shows how closely related certain populations are genetically: Groups which cluster closely are more genetically similar than groups which are further apart. The black star symbol shows where this PGP-UK participant sits in relation to other populations, indicating their ancestry and their most closely related populations according to genetic sequence.

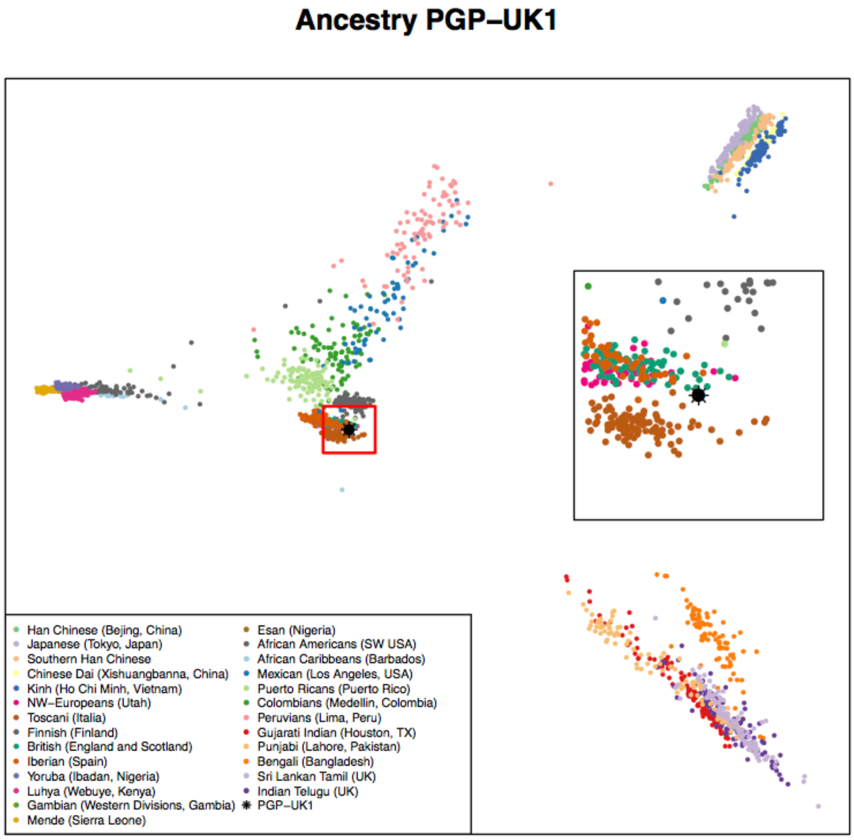


Figure 5: Ancestry Principal Component Analysis

#### 3 Traits (based on SNPedia information)

Existing research has associated many variants with phenotypic traits, some of which can be perceived as beneficial while others appear to have a harmful effect. Some traits are complex and can be affected by several variants. It is likely that some of these would confer a higher risk while others a lower risk of trait manifestation. These can not be combined linearly to produce an actual risk of disease.

It is important to note that in most cases genomic data is probabilistic, not deterministic- i.e. having a genetic predisposition for a disease is not a diagnosis; rather, it shows an increased likelihood of developing that disease. Also, one person can have both potentially beneficial and harmful variants in the same gene, or associated with the same disease.

Some variants can also affect certain populations more, or will only affect a particular gender. For example, a variant for higher risk of endometriosis in the sequence of a male will not directly affect that person, but can be passed on to descendants.

While many traits are the result of a unique variant, many are the combination of several variants throughout the genome. In SNPedia, these are called genosets. These can integrate some of the information already present in the single variant tables, or be the combination of variants that have no phenotypic effect on their own, but contribute to a trait when together.

The variants in the following tables are sorted by magnitude. This is an subjective measure defined in SNPedia to highlight the perceived importance of the genotype described. At the moment this scale goes from 0 to 10. You can read more about it by visiting their explanatory webpage.

As our knowledge grows, the interpretation of the effect of certain variants might change. Clicking on the links in the genome report tables will take you to websites containing more information about each variant.

- Possibly Beneficial Traits

| Mag. | Identifier | Genotype | Summary | ExAC | GetEvidence | ClinVar |
|---|---|---|---|---|---|---|
| 2.5 | rs2943634 | (A;A) | Lower risk of ischemic stroke | | Link | |
| 2.2 | rs2511989 | (A;A) | 0.44x decreased age-related macular degeneratio... | | Link | |
| 2.1 | rs738409 | (G;G) | Most common genotype; slightly less damage from... | Link | Link | |
| 2.1 | rs806380 | (G;G) | Uncommon. lowest odds of cannabis dependence | | | |
| 2 | rs1026732 | (A;A) | <0.70x risk for restless legs | | Link | |
| 2 | rs10503669 | (A;C) | Associated with higher HDL cholesterol | | Link | |
| 2 | rs10504861 | (A;G) | Reduced risk of migraine without aura | | | |
| 2 | rs11635424 | (A;A) | <0.70x risk for restless legs | | Link | |
| 2 | rs12593813 | (A;A) | <0.71x risk for restless legs | | Link | |
| 2 | rs12678919 | (A;G) | Associated with higher HDL cholesterol | | Link | |
| 2 | rs12979860 | (C;C) | ~80% of such hepatitis C patients respond to tr... | | Link | |
| 2 | rs1544410 | (G;G) | Decreased risk of low bone mineral density diso... | | Link | |
| 2 | rs1799884 | (G;G) | Mothers have typical Birth-Weight babies. Sligh... | | | |
| 2 | rs1800972 | (G;G) | Reduced risk for Crohn's disease; reduced risk ... | Link | | |
| 2 | rs1864163 | (A;G) | Associated with higher HDL cholesterol | | Link | |
| 2 | rs2060793 | (A;A) | Lower serum levels of vitamin D | | | |
| 2 | rs2235015 | (G;T) | Somewhat more likely to respond to certain anti... | Link | Link | |
| 2 | rs261332 | (A;G) | Associated with higher HDL cholesterol | | | |
| 2 | rs3738579 | (C;T) | 0.5x decreased risk for cervical cancer: HNSCC:... | | | |
| 2 | rs3764261 | (G;T) | Associated with higher HDL cholesterol | | Link | Link |
| 2 | rs3914132 | (C;T) | Lower otosclerosis risk | | Link | |
| 2 | rs4143094 | (G;G) | No increased risk of colorectal cancer correlat... | | | |
| 2 | rs4149268 | (G;G) | Associated with higher HDL cholesterol | | Link | |
| 2 | rs4585 | (G;G) | Slightly higher (1.35x) odds of good metformin ... | | | |
| 2 | rs505922 | (T;T) | Blood type O | | Link | |
| 2 | rs6505162 | (A;C) | 0.58x decreased risk for esophageal cancer | Link | | |
| 2 | rs6511720 | (G;T) | Slightly lower odds of developing CHD. | | Link | |
| 2 | rs7776725 | (T;T) | Stronger bones | | Link | |

| Mag. | Identifier | Genotype | Summary | ExAC | GetEvidence | ClinVar |
|---|---|---|---|---|---|---|
| 2 | rs801114 | (T;T) | 0.78x decreased Basal Cell Carcinoma risk. | | Link | |
| 1.5 | rs11136000 | (C;T) | 0.84x decreased risk for Alzheimer's disease | | Link | |
| 1.5 | rs309375 | (G;G) | Smaller mosquito bites | | | |
| 1.5 | rs3851179 | (A;A) | 0.85x decreased risk for Alzheimer's disease | | Link | |
| 1.5 | rs4149274 | (C;C) | Associated with higher HDL (good) cholesterol. | | | |
| 1.5 | rs4939883 | (C;T) | Associated with higher HDL cholesterol | | Link | |
| 1.2 | rs11172113 | (C;C) | 0.8x lower risk for migraines | | | |
| 1.2 | rs11246226 | (A;C) | Decreased risk of schizophrenia in limited stud... | | Link | |
| 1.2 | rs6048 | (G;G) | Slightly lower risk (10-20%) of deep vein throm... | Link | Link | Link |
| 1.1 | rs2293347 | (G;G) | Among NSCLC patients: better Gefitinib response... | Link | | |
| 1 | rs10248420 | (A;G) | 7x more likely to respond to certain antidepres... | | Link | |
| 1 | rs11983225 | (C;T) | 7x more likely to respond to certain antidepres... | | Link | |
| 1 | rs182549 | (C;T) | Can digest milk. | | | Link |
| 1 | rs2235040 | (A;G) | 7x more likely to respond to certain antidepres... | Link | Link | |
| 1 | rs2235067 | (A;G) | 7x more likely to respond to certain antidepres... | | | |
| 1 | rs2494732 | (T;T) | Lower odds of psychosis | Link | Link | |
| 1 | rs2952768 | (C;T) | Slightly less drug dependence: decreased effect... | | | |
| 1 | rs4148739 | (A;G) | 7x more likely to respond to certain antidepres... | | Link | |
| 1 | rs4939827 | (C;T) | 0.86x decreased risk for colorectal cancer | | Link | |
| 1 | rs7850258 | (A;G) | Typical odds of developing primary hypothyroidi... | | | |
| 1 | rs800292 | (C;T) | 1% decreased risk of macular degeneration | Link | Link | Link |
| 0.1 | rs891512 | (G;G) | Lower blood pressure than those with an A allel... | Link | | |
| 0 | rs1047781 | (A;A) | ABH blood group "Secretor" status if Japanese | Link | Link | Link |
| 0 | rs10897346 | (C;C) | If depressed: 2.6x more likely to not respond t... | | | |
| 0 | rs1126742 | (T;T) | Higher hypertension risk | Link | Link | |
| 0 | rs12252 | (T;T) | More resistant to influenza | Link | | Link |
| 0 | rs12593929 | (A;A) | Blue eye color more likely | | | |
| 0 | rs16990018 | (A;A) | PrP Codon 171 Asn - Non-pathogenic variant | Link | | Link |
| 0 | rs17244841 | (A;A) | More responsive to statin treatment | | Link | |
| 0 | rs1799782 | (C;C) | Lower risk for skin cancer | Link | Link | |
| 0 | rs1799945 | (C;C) | Not a H63D hemochromatosis carrier. | Link | Link | Link |
| 0 | rs1800562 | (G;G) | Not a C282Y hemochromatosis carrier. | Link | Link | Link |
| 0 | rs2240203 | (A;A) | Blue eye color more likely | | | |
| 0 | rs2306402 | (C;C) | 1.18x increased risk for LOAD | | | |
| 0 | rs28933385 | (G;G) | Prion protein Codon 200 (E) - Non pathogenic va... | | | Link |
| 0 | rs403016 | (C;C) | 2x risk for lupus | | Link | |
| 0 | rs5746059 | (A;A) | Slightly higher fat mass | | | |
| 0 | rs6259 | (G;G) | Best inverse correlation between tea-drinking: ... | Link | Link | |
| 0 | rs74315403 | (G;G) | PrP codon 178 (D) - non pathogenic variant | | | Link |
| 0 | rs7495174 | (A;A) | Blue/gray eyes more likely | | Link | |
| 0 | rs8028689 | (T;T) | Blue eye color if part of blue eye color haplot... | | | |

- Possibly Harmful Traits

| Mag. | Identifier | Genotype | Summary | ExAC | GetEvidence | ClinVar |
|---|---|---|---|---|---|---|
| 3 | rs2066847 | (-;C) | 3x higher risk of Crohn's disease | Link | | Link |
| 3 | rs2981582 | (C;T) | 1.3x higher risk of ER+ breast cancer | | Link | |
| 2.7 | rs10830963 | (C;G) | Increased type-2 diabetes risk; higher gestatio... | | Link | |
| 2.5 | rs1121980 | (C;T) | 1.67x risk for obesity | | Link | |
| 2.5 | rs12803066 | (A;G) | Increased risk of myopia | | | |
| 2.5 | rs13266634 | (C;T) | Increased risk for type-2 diabetes | Link | Link | Link |
| 2.5 | rs1421085 | (C;T) | ~1.3x increased obesity risk | | Link | Link |
| 2.5 | rs16969968 | (A;G) | Slightly higher risk for nicotine dependence: l... | Link | Link | Link |
| 2.5 | rs2254958 | (C;C) | 1.61x increased risk for Alzheimer's | | | |
| 2.5 | rs3738919 | (C;C) | 1.94x risk of developing rheumatoid arthritis | | | |
| 2.5 | rs3780374 | (A;G) | Substantially increased odds of developing V617... | | | |
| 2.5 | rs5888 | (C;T) | 3x higher risk for age-related macular degenera... | Link | | |
| 2.5 | rs664143 | (T;T) | Higher risk for number of cancers | | | |
| 2.5 | rs8034191 | (C;T) | 1.27x lung cancer risk | | Link | |
| 2.3 | rs1859962 | (G;G) | 1.28x increased risk for prostate cancer | | Link | |
| 2.3 | rs7966230 | (C;G) | Slightly lower levels of plasma VWF | | | |
| 2.2 | rs2231137 | (G;G) | ~1.5-3x increased risk for ischemic stroke | Link | Link | Link |
| 2.1 | rs10427255 | (C;C) | Highest odds of photic sneeze reflex | | | |
| 2.1 | rs10811661 | (T;T) | 1.2x increased risk for type-2 diabetes | | Link | |
| 2.1 | rs17070145 | (C;C) | Reduced memory abilities | | | Link |
| 2.1 | rs17563 | (C;C) | Risk for otosclerosis | Link | Link | Link |
| 2.1 | rs4430796 | (A;A) | 1.38x increased risk for prostate cancer | | Link | |
| 2.1 | rs4444903 | (G;G) | 3.5x risk of hep-cancer in cirrhosis patients; ... | | | |
| 2.1 | rs646776 | (A;A) | 1.2x risk of coronary artery disease | | Link | |
| 2.1 | rs7837688 | (G;G) | 1.7x increased risk for prostate cancer | | | |
| 2.1 | rs795484 | (A;G) | Increased morphine dose requirement and postope... | | | |
| 2.1 | rs944289 | (C;T) | 1.3x increased thyroid cancer risk | | Link | |
| 2 | rs10086908 | (C;T) | 1.7x increased risk for prostate cancer | | | |
| 2 | rs1045642 | (C;T) | Slower metaboliser for some drugs | Link | Link | |
| 2 | rs10488631 | (C;T) | 2x increased risk of developing SLE; 1.6x incre... | | Link | |
| 2 | rs1050631 | (C;T) | Mean Survival Time of 25 months for esophageal ... | Link | | |
| 2 | rs1051730 | (C;T) | 1.3x increased risk of lung cancer | Link | Link | Link |
| 2 | rs10871777 | (A;G) | Adults likely to be 0.22 BMI units higher | | | |
| 2 | rs10889677 | (A;C) | 1.5x increased risk for certain autoimmune dise... | | Link | |
| 2 | rs10984447 | (A;A) | >1.17x increased risk for multiple sclerosis | | Link | |
| 2 | rs11045585 | (A;G) | 63% chance (higher than average) of docetaxel-i... | | Link | |
| 2 | rs11190870 | (C;T) | Possibly increased risk of scoliosis | | | |
| 2 | rs1160312 | (A;G) | 1.6x increased risk of Male Pattern Baldness. | | Link | |
| 2 | rs1219648 | (A;G) | 1.20x risk for breast cancer | | Link | |
| 2 | rs12567232 | (A;G) | Increased risk for Crohn's Disease | | Link | |
| 2 | rs13254738 | (A;C) | 1.18x prostate cancer risk | | Link | |
| 2 | rs1333048 | (A;C) | 1.3x increased coronary artery disease risk | | | |
| 2 | rs13376333 | (T;T) | ~2x higher risk of atrial fibrillation | | Link | |
| 2 | rs1360780 | (C;T) | 1.3x increased risk for depression | | Link | |
| 2 | rs144848 | (G;G) | Very slightly increased breast cancer risk | Link | Link | Link |
| 2 | rs1585215 | (A;G) | 2x increased risk for Hodgkin lymphoma | | | |
| 2 | rs1691053 | (A;G) | Increased risk of developing prostate cancer | | | |
| 2 | rs16942 | (A;G) | Very slightly increased breast cancer risk | Link | Link | Link |
| 2 | rs16944 | (G;G) | Increased risk of mental disorders | | Link | |
| 2 | rs1734791 | (A;A) | 1.4x increased risk for lupus | | | |
| 2 | rs17782313 | (C;T) | Adults likely to be 0.22 BMI units higher | | Link | |
| 2 | rs2073963 | (G;T) | Increased risk of baldness | | | |
| 2 | rs2143340 | (C;T) | Increased risk of dyslexia and poor reading per... | | | |

| Mag. | Identifier | Genotype | Summary | ExAC | GetEvidence | ClinVar |
|---|---|---|---|---|---|---|
| 2 | rs2201841 | (C;T) | 1.5x increased risk for Crohn's disease; 2x inc... | | Link | |
| 2 | rs2230201 | (G;G) | >1.4x risk of lupus | Link | | |
| 2 | rs2274223 | (A;G) | 1.5x increased risk for stomach and esophageal ... | Link | Link | |
| 2 | rs2383206 | (A;G) | 1.4x increased risk for heart disease | | | |
| 2 | rs2383207 | (A;G) | Increased risk for heart disease | | | |
| 2 | rs241448 | (C;T) | 1.51x increased risk for Alzheimer's | Link | | Link |
| 2 | rs2420946 | (C;T) | 1.20x risk for breast cancer | | | |
| 2 | rs25487 | (G;G) | 2x higher risk for skin cancer; possibly other ... | Link | Link | |
| 2 | rs2707466 | (G;G) | Weaker bones | Link | Link | |
| 2 | rs2736990 | (C;C) | Increased risk of developing Parkinson's Diseas... | | Link | |
| 2 | rs27388 | (A;A) | Increased risk of developing schizophrenia | | | |
| 2 | rs2908004 | (C;C) | Weaker bones | Link | Link | |
| 2 | rs3212227 | (A;C) | Significantly increased risk of developing cerv... | | | |
| 2 | rs358806 | (C;C) | 1.78x increased risk of developing Type-2 diabe... | | Link | |
| 2 | rs3775948 | (G;G) | Slightly higher risk for gout | | | |
| 2 | rs4027132 | (A;A) | 1.51x increased risk of developing bipolar diso... | | | |
| 2 | rs4129148 | (C;G) | 3x risk of schizophrenia. | | Link | |
| 2 | rs4633 | (T;T) | Higher risk for endometrial cancer | Link | Link | |
| 2 | rs4792311 | (A;G) | Increased risk of prostate cancer | Link | Link | Link |
| 2 | rs493258 | (G;G) | 1.15x risk of Age Related Macular Degeneration | | | |
| 2 | rs4968451 | (A;C) | 1.61x increased risk for meningioma | | | |
| 2 | rs520354 | (A;A) | Increased risk in men for biliary conditions | | | |
| 2 | rs5759167 | (T;T) | Higher prostate cancer risk | | Link | |
| 2 | rs6441286 | (G;T) | 1.54x chance of developing primary biliary cirr... | | Link | |
| 2 | rs6457617 | (C;T) | 2.3x risk of rheumatoid arthritis | | Link | |
| 2 | rs6603272 | (G;T) | 2.74x increased risk of developing schizophreni... | | | |
| 2 | rs6896702 | (T;T) | Increased risk of developing Parkinson's Diseas... | | | |
| 2 | rs6897932 | (C;T) | 1.3x increased risk for multiple sclerosis | Link | Link | Link |
| 2 | rs6922269 | (A;A) | 1.6x risk of coronary artery disease | | Link | |
| 2 | rs6997709 | (G;T) | 1.2x higher risk for hypertension | | | |
| 2 | rs699 | (C;T) | Increased risk of hypertension | Link | Link | Link |
| 2 | rs7216389 | (T;T) | 1.5x increased risk for Childhood Asthma. | | Link | |
| 2 | rs7442295 | (A;A) | ~4x higher risk for hyperuracemia | | Link | |
| 2 | rs744373 | (C;T) | 1.17x risk of Alzheimer's | | | |
| 2 | rs7536563 | (A;A) | >1.12x risk of multiple sclerosis | | Link | |
| 2 | rs7794745 | (A;T) | Slightly increased risk for autism | | Link | Link |
| 2 | rs7807268 | (C;G) | 1.3x risk for Crohn's disease | | Link | |
| 2 | rs7961152 | (A;C) | 1.2x higher risk for hypertension | | | |
| 2 | rs828907 | (G;T) | Slightly increased risk of bladder cancer and 2... | | | |
| 2 | rs9652490 | (A;A) | ~2x increased risk for Parkinson's disease: and... | | Link | |
| 2 | rs965513 | (A;G) | 1.7x increased thyroid cancer risk | | Link | |
| 2.0 | rs2156921 | (G;G) | 1.29x increased risk for depression | | | |
| 2.0 | rs2305795 | (A;A) | 1.64x higher risk of narcolepsy compared to (G;... | | | Link |
| 2.0 | rs9642880 | (T;T) | 1.5x increased bladder cancer risk | | Link | |
| 1.7 | rs8055236 | (G;T) | 1.9x risk for heart disease | | Link | |
| 1.6 | rs1537415 | (C;G) | 1.6x increased risk for periodontitis | | Link | |
| 1.6 | rs2046210 | (T;T) | 1.6x increased breast cancer risk in certain wo... | | Link | |
| 1.6 | rs2736100 | (G;G) | 1.6x higher risk for glioma development | | Link | |
| 1.6 | rs356219 | (G;G) | 1.6x increased risk for Parkinson's disease | | | |
| 1.6 | rs3764880 | (A;A) | 1.2 - 1.8x increased tuberculosis risk | Link | Link | |
| 1.5 | rs10492519 | (A;G) | Slightly increased risk of developing prostate ... | | | |
| 1.5 | rs10757272 | (C;T) | 1.30x increased risk for Coronary artery diseas... | | | |
| 1.5 | rs10859871 | (A;C) | Slight (~1.2x) increase in endometriosis risk | | | |
| 1.5 | rs10883365 | (A;G) | 1.2x increased risk for developing Crohn's dise... | | Link | |
| 1.5 | rs12037606 | (A;G) | 1.22x risk of developing Crohn's disease | | | |

| Mag. | Identifier | Genotype | Summary | ExAC | GetEvidence | ClinVar |
|---|---|---|---|---|---|---|
| 1.5 | rs12210050 | (C;T) | Slightly higher risk for basal cell carcinoma | | Link | |
| 1.5 | rs1223271 | (A;G) | Slightly increased risk of developing Parkinson... | | Link | |
| 1.5 | rs12431733 | (C;T) | Slightly increased risk of developing Parkinson... | | Link | |
| 1.5 | rs12498742 | (A;A) | 1.25 increased risk for gout | | | |
| 1.5 | rs13149290 | (C;C) | Slightly increased risk of developing prostate ... | | | |
| 1.5 | rs1375144 | (C;T) | 1.32x increased risk of developing bipolar diso... | | | |
| 1.5 | rs140701 | (A;A) | Increased risk for anxiety disorders | | | |
| 1.5 | rs17221417 | (C;G) | 1.3x higher risk for Crohn's disease | | Link | |
| 1.5 | rs1801274 | (T;T) | Complex; generally greater risk for cancer prog... | Link | Link | Link |
| 1.5 | rs2272127 | (C;C) | Associated with herpes and schizophrenia | | | |
| 1.5 | rs2280714 | (A;A) | 1.4x increased risk of SLE | | | |
| 1.5 | rs28694718 | (A;A) | >2x higher risk for schizophrenia | | | |
| 1.5 | rs3087243 | (A;G) | Increased risk for auto-immune diseases | | Link | |
| 1.5 | rs356220 | (T;T) | Increased risk of Parkinson's Disease | | | |
| 1.5 | rs3745516 | (A;G) | Slightly increased risk of developing primary b... | | | |
| 1.5 | rs3814570 | (C;T) | 1.3x increased risk for Crohn's disease with il... | | | |
| 1.5 | rs3825776 | (A;G) | 1.3x increased risk for ALS | | Link | |
| 1.5 | rs393152 | (A;A) | Increased risk of both PD and AD | Link | Link | |
| 1.5 | rs401681 | (C;T) | ~1.2x increased risk for several types of cance... | | Link | |
| 1.5 | rs4464148 | (C;T) | 1.10x increased risk for colorectal cancer | | | |
| 1.5 | rs4538475 | (A;G) | Slightly increased risk of developing Parkinson... | | Link | |
| 1.5 | rs4626664 | (A;G) | 1.44x increased risk of developing restless leg... | | Link | |
| 1.5 | rs464049 | (T;T) | Increased risk of schizophrenia in limited stud... | | | |
| 1.5 | rs486907 | (A;G) | 1.5x increased prostate cancer risk | Link | Link | Link |
| 1.5 | rs4982731 | (C;C) | Possible higher risk of childhood acute lymphob... | | | |
| 1.5 | rs5219 | (C;T) | 1.3x increased risk for type-2 diabetes | Link | Link | Link |
| 1.5 | rs6435862 | (G;T) | 1.7x higher risk of aggressive neuroblastoma | | Link | |
| 1.5 | rs6498169 | (A;G) | 1.14x risk of multiple sclerosis | | Link | |
| 1.5 | rs6601764 | (C;T) | 1.16x increased risk of developing Crohn's dise... | | Link | |
| 1.5 | rs6908425 | (C;T) | 1.63x increased risk of developing Crohn's dise... | | Link | |
| 1.5 | rs699473 | (C;T) | ~1.5x increased brain tumor risk | | | |
| 1.5 | rs700651 | (A;G) | ~1.18x increased risk of aneurysm | | Link | |
| 1.5 | rs7774434 | (C;T) | Slightly increased risk of developing primary b... | | | |
| 1.5 | rs9561778 | (G;T) | ~2x increased risk of adverse drug reactions fr... | | Link | |
| 1.5 | rs966221 | (C;C) | 1.5x increased stroke risk certain populations | | | |
| 1.5 | rs995030 | (G;G) | Non-protective against testicular cancer | | Link | |
| 1.4 | rs3131296 | (G;G) | 1.4x increased risk for schizophrenia | | Link | |
| 1.4 | rs4795067 | (G;G) | Slight increase in risk for psoriatic arthritis... | | | |
| 1.4 | rs6010620 | (G;G) | 1.4x higher risk for glioma development; but th... | | Link | |
| 1.3 | rs1042713 | (A;G) | 1.3x increased risk that pediatric inhaler use ... | Link | Link | Link |
| 1.3 | rs10947262 | (C;C) | 1.3x increased risk for osteoarthritis | | | |
| 1.3 | rs110419 | (A;G) | 1.3x increased risk for neuroblastoma | | | |
| 1.3 | rs1260326 | (C;T) | Slightly higher risk for gout | Link | Link | Link |
| 1.3 | rs13361189 | (C;T) | 1.3x increased risk for Crohn's disease | | Link | |
| 1.3 | rs1434536 | (A;G) | 1.29x increased breast cancer risk | | | |
| 1.3 | rs34330 | (C;T) | 1.3x higher risk for endometrial cancer (in Chi... | | | |
| 1.3 | rs4958847 | (A;G) | 1.3x increased risk for Crohn's disease | | | |
| 1.25 | rs13387042 | (A;A) | 1.24x increased risk for breast cancer | | Link | |
| 1.2 | rs10865331 | (A;G) | 1.2x higher risk for ankylosing spondylitis | | | |
| 1.2 | rs1344706 | (T;T) | 1.2x increased risk for schizophrenia | | Link | |
| 1.2 | rs143383 | (C;T) | 1.1x increased risk for osteoarthritis | | Link | Link |
| 1.2 | rs1800693 | (A;G) | Slight (1.2x) increase in risk for multiple scl... | Link | Link | Link |
| 1.2 | rs2056116 | (A;G) | 1.18x risk for breast cancer | | | |
| 1.2 | rs2072590 | (G;T) | 1.2x increased risk for ovarian cancer | | | |
| 1.2 | rs2076295 | (G;T) | One copy of the risk allele (G): slightly incre... | | | |

| Mag. | Identifier | Genotype | Summary | ExAC | GetEvidence | ClinVar |
|---|---|---|---|---|---|---|
| 1.2 | rs2252586 | (A;G) | 1.2x higher risk for glioma development | | | |
| 1.2 | rs6897876 | (C;C) | Slight increase in testicular cancer risk for m... | | | |
| 1.2 | rs8050136 | (A;C) | 1.2x increased risk for T2D in some populations... | | Link | |
| 1.17 | rs17465637 | (A;C) | 1.17x higher risk for myocardial infarction | Link | Link | |
| 1.15 | rs748404 | (C;T) | Very slightly increased risk (1.15) for lung ca... | | Link | |
| 1.1 | rs11037909 | (C;T) | 1.27x type II diabetes risk | Link | | |
| 1.1 | rs11110912 | (C;G) | 1.3x high blood pressure risk | | | |
| 1.1 | rs1799966 | (A;G) | Very slightly increased breast cancer risk | Link | Link | Link |
| 1.1 | rs249954 | (C;T) | Slight if any increased risk of Breast Cancer | | | Link |
| 1.1 | rs2651899 | (A;G) | 1.1x higher risk for migraines | | | |
| 1.1 | rs2653349 | (G;G) | 2-6x increased risk for cluster headaches | Link | Link | |
| 1.1 | rs34516635 | (G;G) | Less longevity for Ashkenazi Jewish women. | Link | | Link |
| 1.1 | rs3740878 | (A;G) | 1.26x type II diabetes risk | Link | | |
| 1.1 | rs3818361 | (C;T) | 1.15x increased risk for late-onset Alzheimer's... | | | |
| 1.1 | rs4977574 | (A;G) | Some studies - but not others - report a slight... | | Link | |
| 1.1 | rs7412 | (C;T) | More likely to gain weight if taking olanzapine... | Link | Link | Link |
| 1.1 | rs925391 | (C;C) | More likely to go bald; common | | | |
| 1.05 | rs2291834 | (C;T) | Very slightly higher risk for myocardial infarc... | | | |
| 1 | rs10761659 | (A;G) | 1.2x risk of Crohn's disease | | Link | |
| 1 | rs2273697 | (A;G) | Adverse reaction more likely to carbamazepine i... | Link | Link | |
| 1 | rs2282679 | (A;C) | Somewhat lower vitamin D levels | | | |
| 1 | rs2546890 | (A;G) | Higher risk of multiple sclerosis | | | |
| 1 | rs3194051 | (A;A) | >1.1x risk of type-1 diabetes | Link | Link | Link |
| 1 | rs6932590 | (T;T) | 1.1x increased risk for schizophrenia | | Link | |
| 1 | rs6974491 | (A;G) | Higher risk of coeliac and/or inflammatory bowe... | | | |
| 1 | rs987525 | (A;C) | 2.5x increased risk for cleft lip | | Link | |
| 0.1 | rs601338 | (A;G) | Susceptible to Norovirus infections | Link | Link | Link |
| 0 | rs1333040 | (C;T) | 1.24x increased myocardial infarction risk: 1.2... | | Link | |
| 0 | rs1800860 | (A;A) | 10% smaller kidneys as newborns | Link | | Link |
| 0 | rs3761418 | (A;A) | 1.3x increased risk for depression | | | |
| 0 | rs4293393 | (T;T) | 1.25x Increased Risk of CKD for T allele in ... | | | |
| 0 | rs440446 | (G;G) | Increased risk in men for biliary conditions | Link | | |
| 0 | rs4714156 | (C;C) | <0.61x risk for restless legs | | | |
| 0 | rs6314 | (C;C) | Higher risk for RA | Link | Link | |

- Genosets (Multi-variant Phenotypes)

| Magnitude | Identifier | Summary |
|---|---|---|
| 4 | gs144 | Male |
| 3.5 | gs126 | Poor warfarin metabolizer |
| 3.3 | gs162 | CYP2C9 Poor Metabolizers |
| 3.1 | gs122 | 7x risk of baldness |
| 3.1 | gs191 | Problem metabolizing NSAIDs |
| 3 | gs241 | Lighter green: brown or hazel eye color |
| 3 | gs273 | Lowest risk (13% of white women) of Atrial Fibr... |
| 2.5 | gs155 | CYP3A5 non-expressor |
| 2.5 | gs157 | More stimulated by coffee |
| 2.5 | gs259 | Homozygous for eye color haplotype #3 |
| 2.5 | gs281 | Part of the 88% of the population claimed not t... |
| 2.5 | gs285 | You will lose 2.5x as much weight on a low fat ... |
| 2.3 | gs255 | Homozygous eye color haplotype #1 |
| 2.1 | gs223 | One copy of GCH1 variant associated with lower ... |
| 2 | gs101 | Probably able to digest milk |
| 2 | gs140 | NAT2 slow metabolizer |
| 2 | gs154 | NAT2 Slow metabolizer |
| 2 | gs173 | CYP2D6*10 |
| 2 | gs221 | Autoimmune disorder risk in Europeans |
| 2 | gs269 | APOE E2/E3 |
| 2 | gs279 | Mild trimethylaminuria |
| 1.5 | gs185 | The beta blocker metoprolol is effective with 1... |
| 1.5 | gs220 | HLA-B*1502? |
| 1.5 | gs247 | Parkinson's Disease Risk |
| 1.2 | gs184 | Able to taste bitterness. |
| 1 | gs182 | CYP2D6*39 |
| 0.1 | gs233 | Normal pain sensitivity |

#### 4 Report Metadata

| Resource | Version | Website |
|---|---|---|
| Genome | GRCh37 | Link |
| BWA | 0.7.12 | Link |
| SAMtools | 1.2 | Link |
| GATK | 3.4-46 | Link |
| PLINK | v1.90b3.35 | Link |
| VEP | 84 | Link |
| SNPedia | 8-Apr-2016 | Link |
| ExAC | v0.3.1 | Link |
| GetEvidence | 8-Apr-2016 | Link |
| ClinVar | 4-Apr-2016 | Link |

Table 5: Analysis Pipeline Versions

Report generated on July 20, 2016 (using report generator version 16-174).

**Additional file 2.** Distribution of private SNVs following effect prediction with multiple methods. SNVs that passed the significance threshold for each method are coloured red.

**A)** CADD

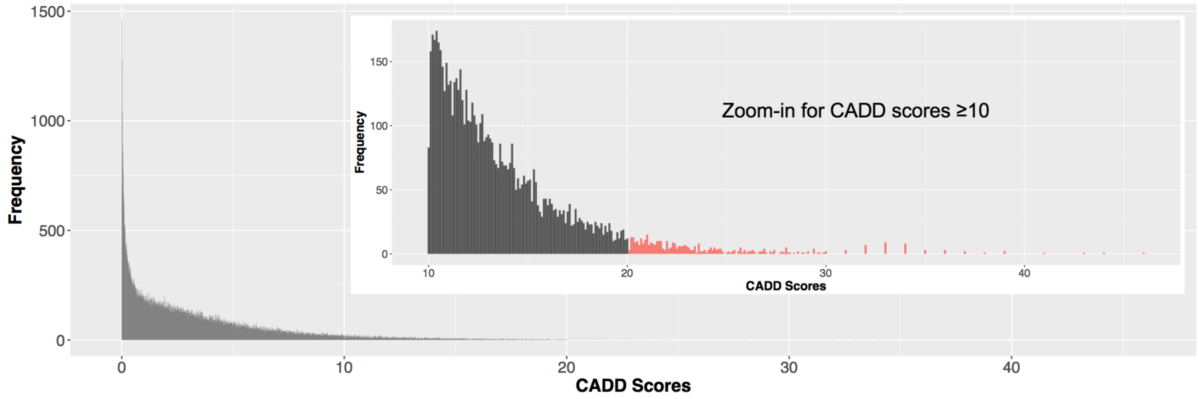


**B)** DANN

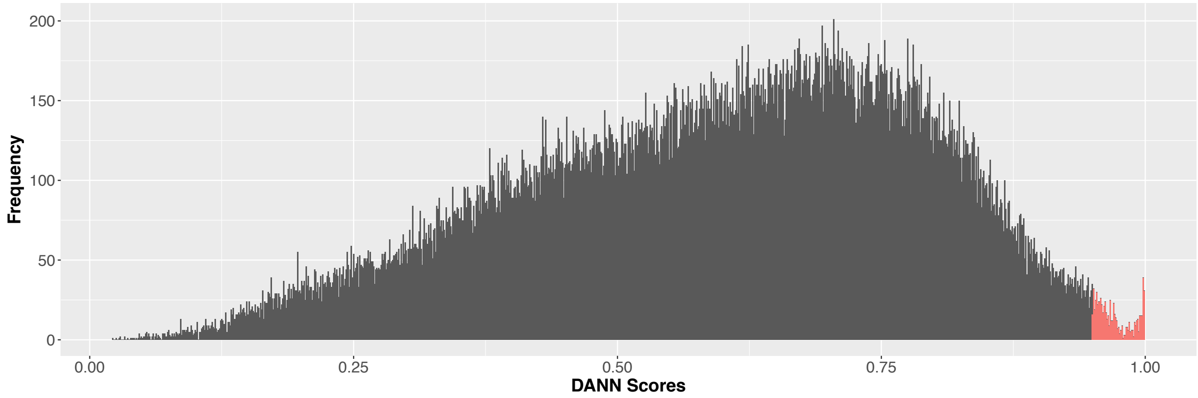


**C)** FATHMM-MKL

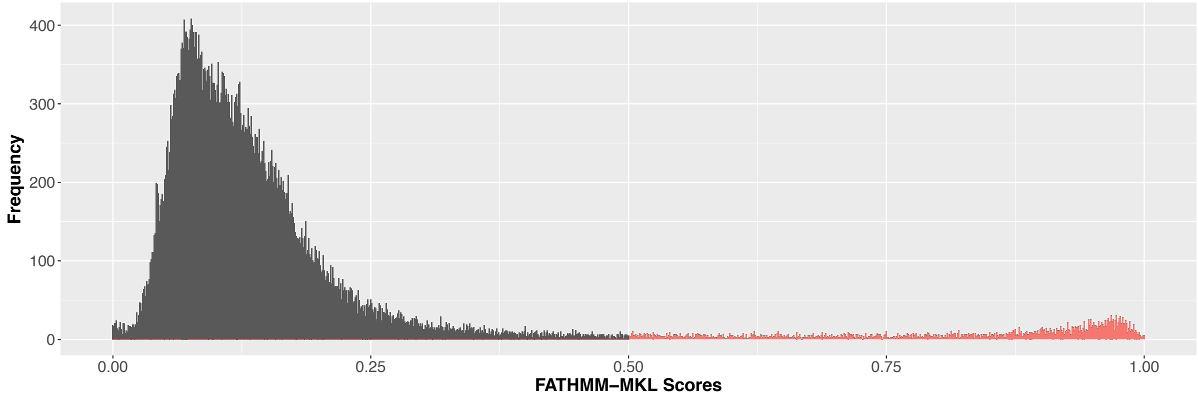

**D)** ExAC-pLI

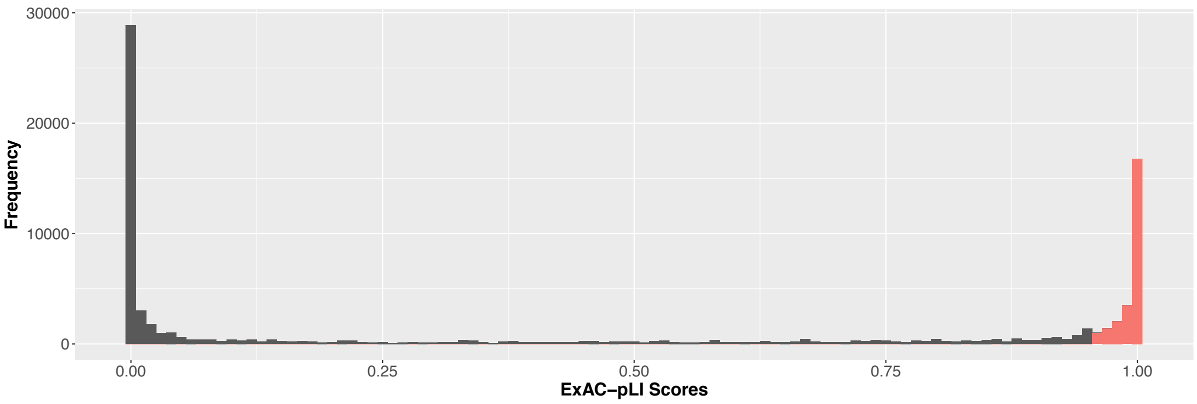
30000
20000
10000
0
Frequency
0.00
0.25
0.50
0.75
1.00
ExAC−pLI Scores

**Additional file 3.** Exemplar PGP-UK 450K methylome report of participant uk35C650 providing predictions for sex, age and smoking status using blood or saliva as indicated.

### Methylome (450K) Report for uk35C650

#### 1 Summary

Epigenetics is the study of modifications of the DNA which control if a gene is switched on or off, without changing the DNA sequence itself. Epigenetic changes are important in many biological processes in human health and disease. There are several different types of epigenetic modifications, of which DNA methylation is the most studied. DNA methylation involves the addition or removal of a methyl group ($CH_3$) to/from cytosine bases in the DNA.

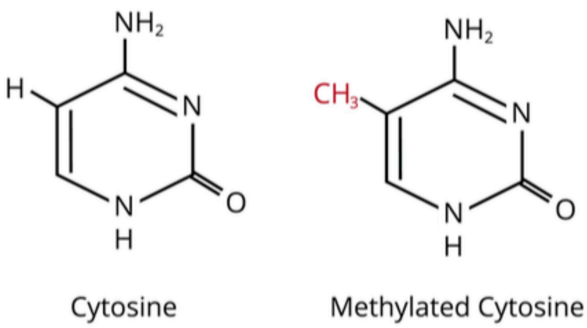


Collectively, all DNA methylation variation within a cell is known as the methylome. The methylome is known to change during normal development, ageing and disease as well as in response to the environment (for example, smoking). It therefore changes throughout life. The methylome is also different in different tissues of the body, such as the brain, skin or blood.

The methylome can be used to predict many features including a person's age, sex and smoking status (current or past/never). DNA methylation differences are widely expected to become biomarkers for environmental exposures, to be used in early diagnosis of disease and to allow matching of patients to the most appropriate disease therapies. As new reliable biomarkers become established they will also be reported for PGP-UK participants.

This report summarises the analysis results of different features from the methylomes of blood and/or saliva. The data were generated using an array-based method from Illumina. The array allows analysis of DNA methylation at around half a million (450K) sites spread across the methylome.

This report was generated automatically and is not clinically approved. It is provided for personal and research purposes only.

#### 2 Prediction of age

A small number of methylation sites in the methylome change throughout a person's lifetime in a predictable way. This allows DNA methylation data to be used to predict a person's current age. By measuring 353 such sites using a methylation array, we have predicted the age of the participant using saliva and/or blood samples. This was carried out using the epigenetic clock [1] which was developed by Steve Horvath at the University of California. If the predicted methylation age deviates from the self-reported actual age at the time of sampling, we further predict age acceleration (where the methylation age is higher than the actual age) and age deceleration (where the methylation age is lower than the actual age). Acceleration and deceleration are shown if the difference is more than 3.6 years (which is the range of accuracy for the epigenetic clock).

Deviations between actual and methylation age can give an insight into general health. Studies have recently associated extreme methylation age acceleration with certain types of cancer [2] and overall mortality [3], while methylation age deceleration has been associated with longevity [4].

| | |
|---|---|
| PGP Participant | uk35C650 |

| | |
|---|---|
| Sample Tissue | Saliva |
| Predicted Age | 53 years, 5 months |
| Age at Sampling | 59 years, 8 months |

| | |
|---|---|
| Sample Tissue | Blood |
| Predicted Age | 52 years, 9 months |
| Age at Sampling | 59 years, 8 months |

| | |
|---|---|
| Sample Tissue | Saliva |
| Predicted Age | 51 years, 4 months |
| Age at Sampling | 59 years, 8 months |

| | |
|---|---|
| Sample Tissue | Blood |
| Predicted Age | 56 years, 4 months |
| Age at Sampling | 61 years, 11 months |

#### 3 Prediction of sex

Females have two X chromosomes but only require one of them to be active. The other X chromosome is inactivated by DNA methylation and other silencing mechanisms. By measuring DNA methylation levels on the X chromosome, sex can be predicted [5].

| | |
|---|---|
| PGP Participant | uk35C650 |
| Self Reported Sex | Male |

| | |
|---|---|
| Sample Tissue | Saliva |
| Predicted Sex | Male |
| Fraction X Chromosome Methylation | 0.337551761450253 |

| | |
|---|---|
| Sample Tissue | Blood |
| Predicted Sex | Male |
| Fraction X Chromosome Methylation | 0.339751421837792 |

| | |
|---|---|
| Sample Tissue | Saliva |
| Predicted Sex | Male |
| Fraction X Chromosome Methylation | 0.332147528543224 |

| | |
|---|---|
| Sample Tissue | Blood |
| Predicted Sex | Male |
| Fraction X Chromosome Methylation | 0.328610239082086 |

#### 4 Prediction of exposure to smoking

One of the most well validated exposures which alters DNA methylation is exposure to tobacco smoking. Many studies have shown that DNA methylation at hundreds of sites across the genome changes when someone smokes, particularly at a gene called AHRR. Studies have also found that while previous smokers still have traces of methylation differences, the DNA methylation changes associated with smoking gradually change to be more similar to the methylation of people who have never smoked. A recent study found that these methylation sites change much more in the buccal cells (cells from the epithelial lining of the mouth) of smokers compared to blood cells.

The smoking status for PGP-UK participants was predicted from saliva and/or blood using 187 methylation sites which have been found to change in smokers [6]. Using a method previously described [7], the methylation levels at these sites were used to generate a weighted methylation score, which can be used to differentiate between past/never and current smokers. It has been demonstrated that if a participant has a score of more than 17.55 for Europeans, or more than 11.79 for South Asians, they are classified as current smoker. A limitation of this measure is that the smoking scores have not been tested comprehensively in people of different ethnicities, so we do not yet know the exact threshold to define smoking status in different ethnicities.

| | |
|---|---|
| PGP Participant | uk35C650 |
| Self Reported Current Smoker | No |
| Self Reported Past Smoker | Yes |

| | |
|---|---|
| Sample Tissue | Saliva |
| Smoking Score | 1.54594612716571 |
| Smoking Prediction | Past/Never Smoker |

| | |
|---|---|
| Sample Tissue | Blood |
| Smoking Score | -1.60092756075582 |
| Smoking Prediction | Past/Never Smoker |

| | |
|---|---|
| Sample Tissue | Saliva |
| Smoking Score | 1.49212699345117 |
| Smoking Prediction | Past/Never Smoker |

| | |
|---|---|
| Sample Tissue | Blood |
| Smoking Score | -1.60989755334765 |
| Smoking Prediction | Past/Never Smoker |

### 5 Appendix

#### 5.1 Methylation sites used in epigenetic age prediction (n=353)

| | | | | | | |
|---|---|---|---|---|---|---|
| cg00075967 | cg00374717 | cg00864867 | cg00945507 | cg01027739 | cg01353448 | cg01584473 |
| cg01644850 | cg01656216 | cg01873645 | cg01968178 | cg02085507 | cg02154074 | cg02217159 |
| cg02331561 | cg02332492 | cg02364642 | cg02388150 | cg02479575 | cg02489552 | cg02580606 |
| cg02654291 | cg02827112 | cg02972551 | cg03103192 | cg03167275 | cg03270204 | cg03565323 |
| cg03588357 | cg03760483 | cg04084157 | cg04126866 | cg04528819 | cg04836038 | cg05250458 |
| cg05294243 | cg05365729 | cg05675373 | cg05755779 | cg05921699 | cg05960024 | cg06121469 |
| cg06144905 | cg06361108 | cg06462291 | cg06493994 | cg06557358 | cg06738602 | cg06810647 |
| cg06952310 | cg06993413 | cg07285276 | cg07291563 | cg07337598 | cg07455279 | cg07595943 |
| cg08030082 | cg08090772 | cg08124722 | cg08251036 | cg08370996 | cg08413469 | cg08434234 |
| cg08771731 | cg08965235 | cg09019938 | cg09118625 | cg09191327 | cg09418283 | cg09509673 |
| cg09785172 | cg09869858 | cg09885951 | cg10281002 | cg10376763 | cg10377274 | cg10486998 |
| cg10523019 | cg10920957 | cg11932564 | cg12351433 | cg12373771 | cg12768605 | cg12830694 |
| cg12946225 | cg13038560 | cg13216057 | cg13319175 | cg13460409 | cg13682722 | cg13836627 |
| cg13854874 | cg13899108 | cg13975369 | cg14258236 | cg14308452 | cg14329157 | cg14424579 |
| cg14501253 | cg14658362 | cg14723032 | cg14894144 | cg14992253 | cg15341340 | cg15381769 |
| cg15547534 | cg15661409 | cg15974053 | cg15988232 | cg16150435 | cg16241714 | cg16494477 |
| cg16547529 | cg16579101 | cg17063929 | cg17099569 | cg17285325 | cg17408647 | cg17655614 |
| cg17729667 | cg17853587 | cg17960516 | cg18055007 | cg18180783 | cg18440048 | cg18573383 |
| cg18983672 | cg18984151 | cg19008809 | cg19167673 | cg19273182 | cg19305227 | cg19346193 |
| cg19478743 | cg19514928 | cg19692710 | cg19945840 | cg20295671 | cg20305610 | cg20524216 |
| cg20692569 | cg20761322 | cg20795863 | cg20828084 | cg20914508 | cg20947775 | cg20999813 |
| cg21096399 | cg21378206 | cg21460081 | cg21801378 | cg21870884 | cg22006386 | cg22289837 |
| cg22432269 | cg22449114 | cg22679120 | cg22736354 | cg22809047 | cg22901840 | cg22920873 |
| cg23517605 | cg23662675 | cg23941599 | cg24116886 | cg24126851 | cg24254120 | cg24262469 |
| cg24450312 | cg24580001 | cg24834740 | cg25070637 | cg25148589 | cg25505610 | cg25552492 |
| cg25683012 | cg25771195 | cg25781123 | cg26003813 | cg26005082 | cg26045434 | cg26297688 |
| cg26372517 | cg26453588 | cg26620959 | cg26842024 | cg26845300 | cg27092035 | cg27169020 |
| cg27319898 | cg27377450 | cg27413543 | cg27494383 | cg00091693 | cg00168942 | cg00431549 |
| cg00436603 | cg01027805 | cg01234063 | cg01262913 | cg01407797 | cg01459453 | cg01485645 |
| cg01511567 | cg01560871 | cg01570885 | cg01820374 | cg02047577 | cg02071305 | cg02275294 |
| cg02335441 | cg03019000 | cg03286783 | cg03330058 | cg03578041 | cg03682823 | cg03891319 |
| cg03947362 | cg04005032 | cg04094160 | cg04121983 | cg04268405 | cg04431054 | cg04452713 |
| cg04474832 | cg04999691 | cg05442902 | cg05590257 | cg05847778 | cg05903609 | cg06044899 |
| cg06117855 | cg06513075 | cg06688848 | cg06836772 | cg06926735 | cg07158339 | cg07388493 |
| cg07408456 | cg07498421 | cg07663789 | cg07730301 | cg07770222 | cg07849904 | cg08186124 |
| cg08331960 | cg09133026 | cg09441152 | cg09646392 | cg09722397 | cg09722555 | cg09809672 |
| cg10045881 | cg10266490 | cg10345936 | cg10865119 | cg10940099 | cg11025793 | cg11299964 |
| cg11314684 | cg11388238 | cg11653266 | cg12413566 | cg12616277 | cg12941369 | cg12985418 |
| cg13129046 | cg13269407 | cg13302154 | cg13547237 | cg13828047 | cg13931228 | cg14060828 |
| cg14163776 | cg14175438 | cg14408969 | cg14409958 | cg14423778 | cg14597908 | cg14654875 |
| cg14727952 | cg15185286 | cg15262928 | cg15703512 | cg15804973 | cg16034652 | cg16168311 |
| cg16358826 | cg16408394 | cg16419345 | cg16744741 | cg16899442 | cg16984944 | cg17274064 |
| cg17324128 | cg17338403 | cg17589341 | cg17686885 | cg18031008 | cg18139769 | cg18328933 |
| cg18956095 | cg19044674 | cg19046959 | cg19420968 | cg19569684 | cg19706682 | cg19722847 |
| cg19724470 | cg19761273 | cg19853760 | cg20100381 | cg20240860 | cg21211748 | cg21305265 |
| cg21370143 | cg21395782 | cg21950518 | cg22171829 | cg22190114 | cg22197830 | cg22568540 |
| cg22613010 | cg22637507 | cg22947000 | cg23092072 | cg23124451 | cg23180365 | cg23786576 |
| cg24058132 | cg24081819 | cg24471894 | cg24888049 | cg24899750 | cg25101936 | cg25159610 |
| cg25166896 | cg25411725 | cg25564800 | cg25657834 | cg25809905 | cg25928579 | cg26043391 |
| cg26162695 | cg26394940 | cg26456957 | cg26614073 | cg26723847 | cg26824091 | cg27015931 |
| cg27016307 | cg27202708 | cg27544190 | | | | |

#### 5.2 Methylation sites used in smoking prediction (n=187)

| | | | | | | |
|---|---|---|---|---|---|---|
| cg09469355 | cg08884752 | cg12547807 | cg04885881 | cg21393163 | cg21913886 | cg19713429 |
| cg27537125 | cg15542713 | cg24049493 | cg23090529 | cg21140898 | cg19406367 | cg25189904 |
| cg09662411 | cg18146737 | cg12876356 | cg18316974 | cg09935388 | cg11231349 | cg08709672 |
| cg20295214 | cg03547355 | cg17819085 | cg23079012 | cg06635952 | cg26271591 | cg23667432 |
| cg03188382 | cg19713851 | cg27241845 | cg03329539 | cg06644428 | cg05951221 | cg21566642 |
| cg01940273 | cg13193840 | cg17024919 | cg15693572 | cg23480021 | cg03274391 | cg00501876 |
| cg18642234 | cg15417641 | cg00336149 | cg21188533 | cg19859270 | cg02657160 | cg25197194 |
| cg08202836 | cg21121843 | cg19719391 | cg24556382 | cg11554391 | cg17924476 | cg08606254 |
| cg12806681 | cg03991871 | cg23916896 | cg11902777 | cg01899089 | cg05575921 | cg26703534 |
| cg01097768 | cg14817490 | cg25648203 | cg21161138 | cg03604011 | cg24090911 | cg13039251 |
| cg05673882 | cg26908328 | cg16786458 | cg14580211 | cg12513616 | cg01882991 | cg06126421 |
| cg14753356 | cg24859433 | cg15342087 | cg17619755 | cg10807309 | cg15474579 | cg00931843 |
| cg00921574 | cg19717773 | cg02451831 | cg08972170 | cg19089201 | cg22132788 | cg04180046 |
| cg12803068 | cg07826859 | cg03440944 | cg21322436 | cg25949550 | cg11207515 | cg17372101 |
| cg12276019 | cg24540678 | cg13518625 | cg19589396 | cg25305703 | cg12075928 | cg26361535 |
| cg13787850 | cg01692968 | cg13910681 | cg22539182 | cg25953130 | cg27312979 | cg25421530 |
| cg01744331 | cg07123182 | cg16556677 | cg26963277 | cg04039799 | cg09197783 | cg16611234 |
| cg19254163 | cg21611682 | cg14624207 | cg01901332 | cg11660018 | cg23771366 | cg03234777 |
| cg26282236 | cg02583484 | cg04158018 | cg23681440 | cg23126342 | cg25491122 | cg06885459 |
| cg17487894 | cg01731783 | cg22851561 | cg24996979 | cg10919522 | cg13976502 | cg13038618 |
| cg05875421 | cg05284742 | cg06819357 | cg26242531 | cg11730703 | cg01208318 | cg15022400 |
| cg03489965 | cg18335991 | cg00310412 | cg11152412 | cg23161492 | cg05194346 | cg01207684 |
| cg09099830 | cg03155159 | cg00911794 | cg23621097 | cg09858022 | cg19572487 | cg04956244 |
| cg16255816 | cg03373393 | cg25809905 | cg21280392 | cg07465627 | cg02186444 | cg07251887 |
| cg06459104 | cg00073090 | cg15187398 | cg07381806 | cg00835193 | cg03636183 | cg15159987 |
| cg23973524 | cg11902728 | cg22649124 | cg11701312 | cg16201146 | cg07339236 | cg00871610 |
| cg06595162 | cg23110422 | cg22635096 | cg02532700 | cg01127300 | | |

#### 6 Raw Data

The raw data used to create this report has been assigned the identifier E-MTAB-5377 in the ArrayExpress Archive hosted at the European Bioinformatics Institute (EBI).

The dataset can be accessed at: https://www.ebi.ac.uk/arrayexpress/experiments/E-MTAB-5377/

#### 7 References

1) https://labs.genetics.ucla.edu/horvath/dnamage/

2) Lin 2015, Epigenetic ageing signatures are coherently modified in cancer (PLoS Genet.) DOI: 10.1371/journal.pgen.1005334

3) Marioni 2015, DNA methylation age of blood predicts all-cause mortality in later life (Genome Biology) DOI: 10.1186/s13059-015-0584-6

4) McEwen 2017, Differential DNA methylation and lymphocyte proportions in a Costa Rican high longevity region (Epigenetics & Chromatin) DOI: 10.1186/s13072-017-0128-2

5) Fortin 2014, Minfi Tutorial BioC2014 (Bioconductor)
https://www.bioconductor.org/help/course-materials/2014/BioC2014/minfi_BioC2014.pdf

6) Zeilinger 2013, Tobacco smoking leads to extensive genome-wide changes in DNA methylation (Plos One) DOI: 10.1371/journal.pone.0063812

7) Elliott 2014, Differences in smoking associated DNA methylation patterns in South Asians and Europeans (Clin Epigenetics) DOI: 10.1186/1868-7083-6-4

Report generated on February 14, 2018.
